# The little skate genome and the evolutionary emergence of wing-like fin appendages

**DOI:** 10.1101/2022.03.21.485123

**Authors:** Ferdinand Marlétaz, Elisa de la Calle-Mustienes, Rafael D. Acemel, Tetsuya Nakamura, Christina Paliou, Silvia Naranjo, Pedro Manuel Martínez-García, Ildefonso Cases, Victoria A. Sleight, Christine Hirschberger, Marina Marcet-Houben, Dina Navon, Ali Andrescavage, Ksenia Skvortsova, Paul Edward Duckett, Álvaro González-Rajal, Ozren Bogdanovic, Johan H. Gibcus, Liyan Yang, Lourdes Gallardo-Fuentes, Ismael Sospedra, Javier Lopez-Rios, Fabrice Darbellay, Axel Visel, Job Dekker, Neil Shubin, Toni Gabaldón, Juan J. Tena, Darío G. Lupiáñez, Daniel S. Rokhsar, José Luis Gómez-Skarmeta

## Abstract

Skates are cartilaginous fish whose novel body plan features remarkably enlarged wing-like pectoral fins that allow them to thrive in benthic environments. The molecular underpinnings of this unique trait, however, remain elusive. Here we investigate the origin of this phenotypic innovation by developing the little skate *Leucoraja erinacea* as a genomically enabled model. Analysis of a high-quality chromosome-scale genome sequence for the little skate shows that it preserves many ancestral jawed vertebrate features compared with other sequenced genomes, including numerous ancient microchromosomes. Combining genome comparisons with extensive regulatory datasets in developing fins – gene expression, chromatin occupancy and three-dimensional (3D) conformation – we find skate-specific genomic rearrangements that alter the 3D regulatory landscape of genes involved in the planar cell polarity (PCP) pathway. Functional inhibition of PCP signaling resulted in marked reduction of anterior fin size, confirming this pathway as a major contributor of batoid fin morphology. We also identified a fin-specific enhancer that interacts with 3’ *HOX* genes, consistent with the redeployment of *Hox* gene expression in anterior pectoral fins, and confirmed the potential of this element to activate transcription in the anterior fin using zebrafish reporter assays. Our findings underscore the central role of genome reorganizations and regulatory variation in the evolution of phenotypes, shedding light on the molecular origin of an enigmatic trait.

## MAIN

The origin and diversification of vertebrates was accompanied by the appearance of key developmental innovations ^1,2^. Among them, paired appendages show an exquisite diversity of forms and adaptations not only in tetrapods, but also in chondrichthyans (cartilaginous fish) where fin structures are remarkably diverse ^1^. One of the most fascinating examples are the wing-like appendages of skates (**Fig. 1a**), in which the pectoral fins extend anteriorly and fuse with the head. This unique structure creates power for forward propulsion and led to the emergence of novel swimming mechanisms that allowed skates to thrive close to the sea floor ^3^. Transcriptomic analysis on skate developing fins revealed a major reorganization of signaling gradients relative to other vertebrates ^3^. In skate, the redeployment of developmental transcription factors such as 3’ *HOX* genes initiates an anterior signaling center analogous to the posterior Apical Ectodermal Ridge (AER). These changes arose ∼286-221 million years ago (**Fig. 1a**) after the divergence between sharks and skates. Nevertheless, the genomic and regulatory changes underlying these novel expression domains have remained elusive.

**Fig.1:**
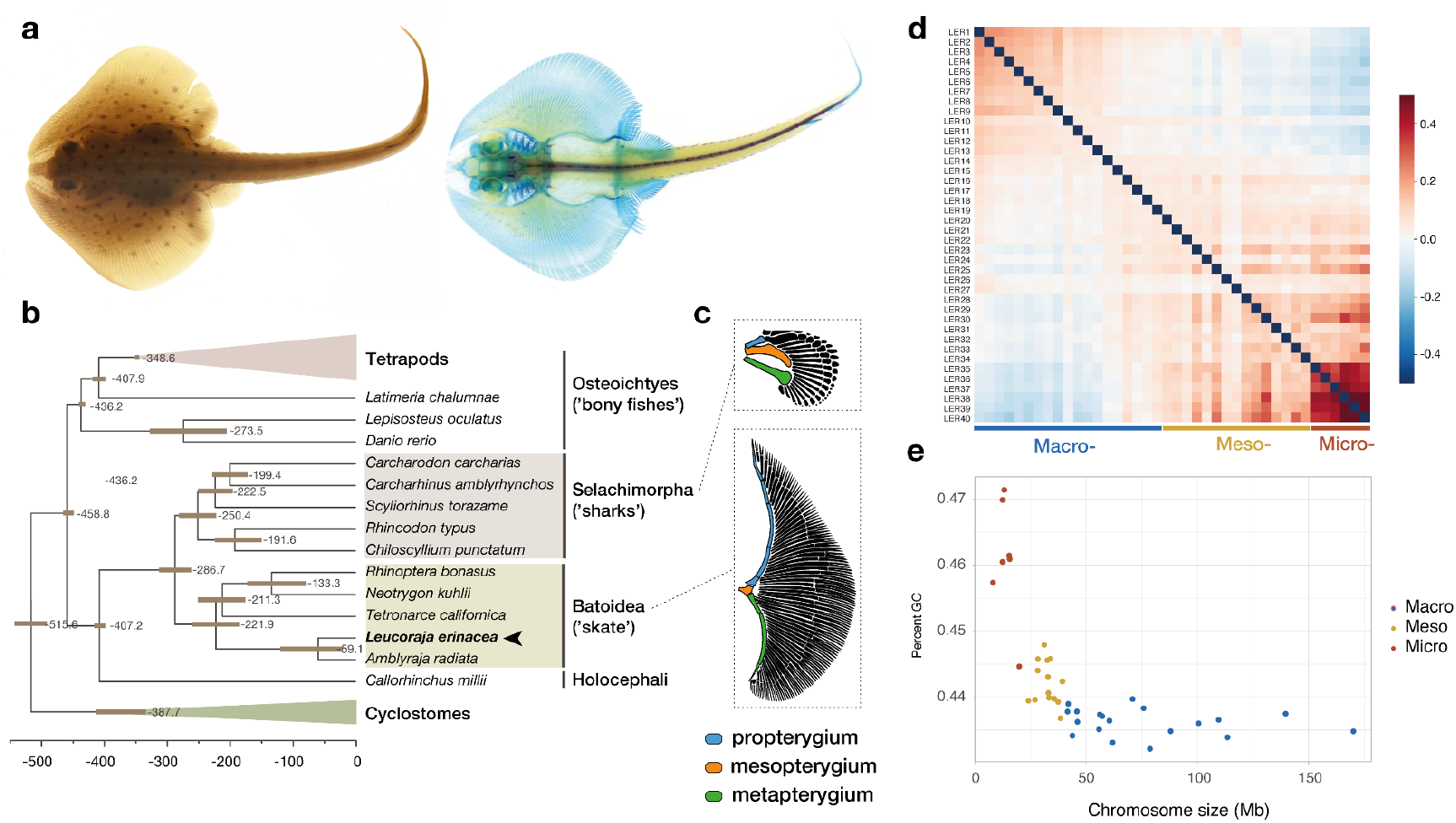
The little skate morphology and genome evolution. **a**, Adult little skate (*Leucoraja erinacea*) and skeletal stainings using alcian blue and alizarin red. **b**, Chronogram showing the branching and divergence time of chondrichthyan and selected osteichthyan lineages (see also **Extended Data Fig. 1**). **c**, Morphological differences of the skeleton between pectoral fins in shark and skate highlighting the expansion of a wing-like fin. **d**, Pairwise Hi-C contact density between 40 skate chromosomes showing an increased interchromosomal interaction between the smallest ones (micro-chromosomes). Color scale: log transformed observed/expected interchromosomal Hi-C contacts. **e**, Little skate chromosomes classification based on the relationship between their size (x-axis) and GC% (y-axis) highlighting the high GC content of microchromosomes.

Many vertebrate evolutionary innovations were influenced by the drastic genomic reorganizations caused by two rounds of whole genome duplication (WGD). The ancestral 18 chordate chromosomes were duplicated, reshuffled and rearranged to give rise to the diversity of existing karyotypes in vertebrates ^4^. Concomitantly, the pervasive loss of paralogous genes after the WGD produced gene deserts enriched in regulatory elements ^5^. Compellingly, those drastic genomic alterations were paralleled by marked changes in gene regulation, contributing to an increase in the pleiotropy of developmental genes ^5^ and on the complexity of their regulatory landscapes ^6^. In vertebrates, regulatory landscapes are spatially organized into topologically associating domains (TADs) ^7,8^. TADs correspond to large genomic regions displaying increased self-interaction frequencies, in which regulatory elements interact with cognate promoters to constitute precise transcriptional patterns. TADs have been consistently found in every vertebrate species where 3D chromatin interactions have been profiled ^9^, suggesting that this mode of spatial genome organization was already present in the gnathostome ancestor. Furthermore, TAD organization influences the evolution of gene order, by directing rearrangement hotspots ^10^. Genomic rearrangements that alter TADs can be a prominent cause of developmental disease or cancer ^11^, but also serve as a source for evolutionary innovation ^12,13^. However, how TAD organization may have impacted the evolution of gene regulation and the subsequent emergence of lineage-specific traits after vertebrate whole genome duplications (WGD) remains largely unexplored.

To gain insight into the evolution of the jawed vertebrate (gnathostome) karyotypes and identify the genomic and regulatory changes underlying the evolution of the skate wing-like appendages, we generated a chromosome-scale assembly of the little skate *Leucoraja erinacea* and performed extensive functional characterization of its developing fins. Our analyses revealed a karyotype configuration resembling that of the gnathostome ancestor, characterized by a slower paralogue loss and smaller chromosomes, which suggests fewer fusion events after the second round (2R) of vertebrate WGD than in other jawed vertebrates. We find that the 3D organization of the skate genome arises from an interplay between transcription-based A/B compartments and TADs likely formed by a loop extrusion mechanism as described for mammals ^14^. The comparison of the 3D organization of the alpha and beta chromosomes after 2R revealed a prominent loss of complete TADs, likely contributing to karyotype stabilization. By further combining RNA-seq and ATAC-seq in anterior and posterior pectoral and pelvic fins we identified the PCP pathway and *HOX* gene regulation as key contributors of batoid fin morphology, which we further validated with functional assays in zebrafish and skate. Our study illustrates how comparative multi-omics approaches can be effectively employed to elucidate the molecular underpinnings of evolutionary traits.

## RESULTS

### Genome sequencing and comparative genomics

We assembled the little skate genome to chromosome scale by integrating long- and short-read genome sequencing data with chromatin conformation capture data (Hi-C). Our assembly includes 40 chromosome-scale (>2.5 Mb) scaffolds including 19 macro chromosomes (>40 Mb), 14 meso chromosomes (between 20 and 40 Mb) and 7 micro chromosomes (<20 Mb) that together represent 91.7% of the 2.2 Gb assembly. This putative chromosome number is within the range reported for other Rajidae species ^15^. Despite technical challenges due to a high level of nucleotide polymorphism (1.6% heterozygosity) and a repeat content dominated by recently expanded LINE retrotransposons (**Extended Data Fig. 2**), our assembly showed a similar or higher degree of completeness with respect to gene content compared with other recently sequenced chondrichthyans (BUSCO, **Extended Data Table 1**).

We annotated 26,715 likely protein-coding genes in the little skate genome using extensive transcriptome resources ^16^, with 23,870 possessing homologs in other species. Using comparative analysis with 20 other sequenced vertebrates we reconstructed the complete set of skate gene evolutionary histories - *i*.*e*., the phylome - and used it to infer patterns of gene duplication and loss, as well as orthology and paralogy relationships (**Extended Data Table 2)**. This rich evolutionary resource for skate genes is available to browse and download at PhylomeDB and MetaPhoRs ^17,18^. We used phylogenomic methods to reconstruct jawed vertebrate phylogeny and infer divergence times, finding a somewhat more ancient divergence between sharks and skates than previously estimated ^19^ (**Fig. 1b, Extended Data Table 2**). Compared with other reported chondrichthyan genomes, *L. erinacea* displays the lowest number of species-specific gene losses (354 losses, see **Extended Data Fig. 1**). Similar to sharks and dogfishes (selachians), ^20,21^ the little skate has larger introns than found in tetrapods (median size 2,167 bp vs 1,586 bp in human) although these introns are not enriched in a particular repeat category.

Skate micro-chromosomes have an overall higher gene density compared than macro-chromosomes (**Extended Data Fig. 2**) which suggests that, as in birds, these small chromosomes are prone to GC-biased gene conversion ^22^. Skate micro-chromosomes also show a high degree of interchromosomal contacts compared with macro-chromosomes (**Fig. 1d, e**), as also found in snakes and other tetrapods ^23^, which suggests this is a general feature of vertebrate micro-chromosomes.

### Chromosome evolution

We surveyed the arrangement of syntenic chromosomal segments derived from ancestral chordate linkage groups (CLGs) in skate, gar, and chicken, using amphioxus as an outgroup as it did not undergo WGDs ^24^, and found that the chromosomal organization of the skate genome closely resembles that of the most recent jawed vertebrate common ancestor (**Fig. 2a**). By classifying single-copy orthologs by their respective genomic locations across multiple species, we assigned ancestral vertebrate chromosomal segments to both rounds of whole genome duplication, dubbed ‘alpha/beta’ and ‘1/2’ ^24^(**Fig. 2b**). The smaller vertebrate chromosomes often show a reciprocal correspondence across multiple species and correspond to a single ancestral gnathostome unit (10 chromosomes have a 1:1:1 orthology bet between skate, gar, and chicken, Figure 2b) ^24–26^. For instance, the trios LER25≡LOC20≡GGA15 and LER28≡LOC22≡GGA19 are both related to a single CLG (CLG-G) and represent the two copies that survived the first genome duplication (‘1R’), while other trios such as LER21≡LOC18≡GGA20 and LER29≡LOC19≡GGA28 are derived from the fusion of several CLGs that is observed in all vertebrate genomes (**Fig. 2b**). The occurrence of these fusions in two traceable copies (on LER4 and LER21) supports a scenario where they fused between an early pan-vertebrate ‘1R’ duplication and a second gnathostome specific ‘2R’ ^24,26^.

**Fig. 2:**
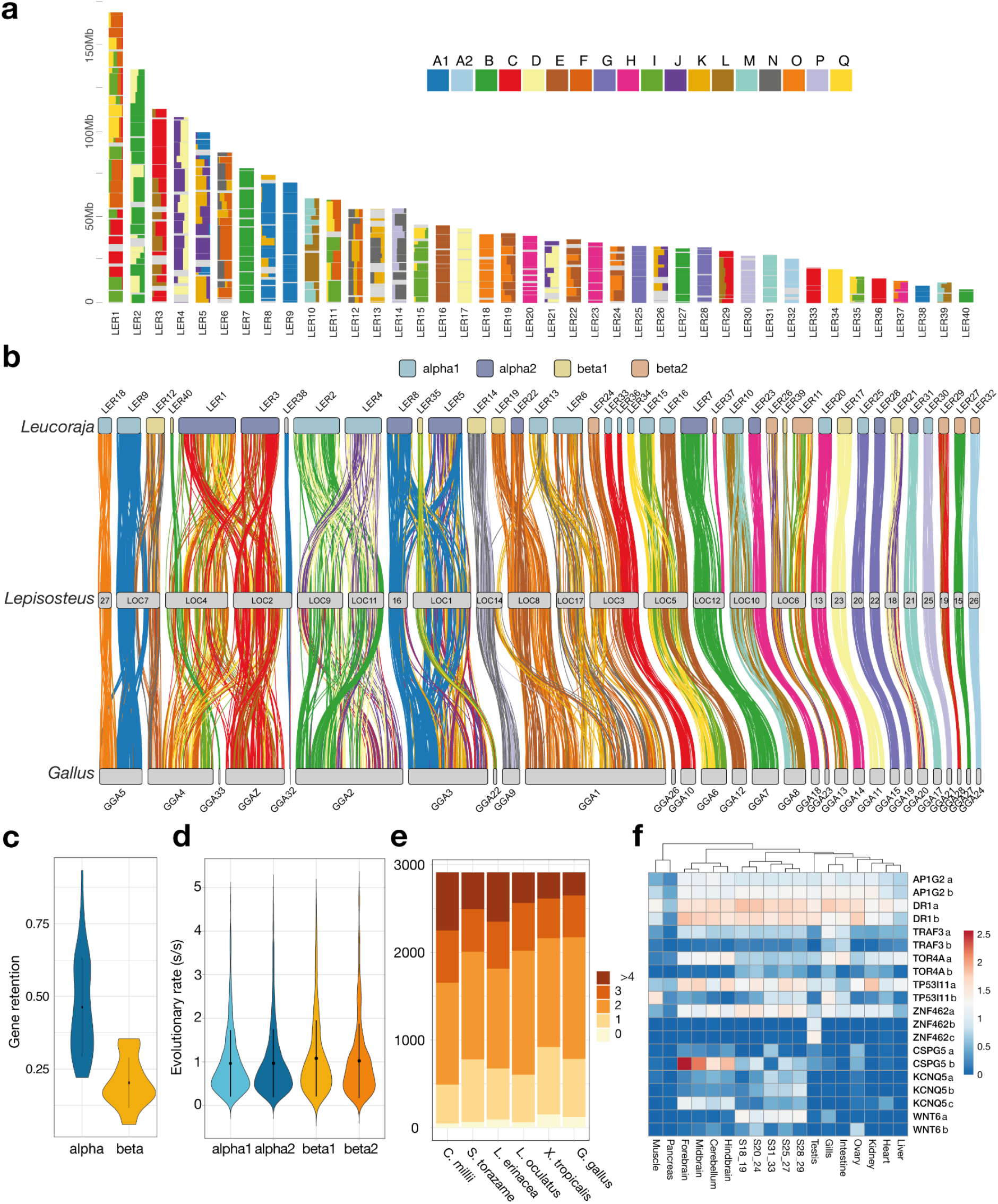
Ancestral linkage and the architecture of early vertebrate genomes. **a**, Fraction of genes derived from each CLG (depicted as squares named A1-Q) in skate chromosomes represented for bins of 20 genes. **b**, Syntenic orthology relationship between skate, gar and chicken relying on genes with a significant ALG assignment in regard to amphioxus. Skate chromosomes are colored by segmental identity and links colored by CLG. **c**, Rates of gene retention for α or β segments derived from the second alloploid event of vertebrate WGD. **d**, Rates of evolution of genes located in α or β segments estimated as ML distance to the amphioxus outgroup (LG+Γ). **e**, Respective gene family composition for ohnologues in selected jawed vertebrates species indicating differential paralogue loss. **f**, Gene expression for selected sets of differentially lost ohnologs for which a copy was lost in the gnathostome but not in the chondrichthyan lineage.

In many vertebrate species, the larger chromosomes derive from fusions of ancestral units (CLGs). We determined that the skate often represents an ancestral state among jawed vertebrates and that subsequent fusions occurred either in chicken (GGA5), in gar (LOC5) or in their common ancestor (LER 2 and 4, see below). For instance, ancestral gnathostome chromosomes resembling skate LER9, LER12, and LER18 fused in different ways to form chromosomes in gar and chicken. Similarly, LER10≡GGA8 and LER23≡GGA18 (≡BFL8) likely represent ancestral units, which fused in gar chromosome LOC10, likely through a centric Robertsonian fusion (**Fig. 2b**). Noticeably, these two chromosomes are also preserved in their ancestral condition in the bowfin, the sister group of gar, implying that this fusion occurred specifically in the gar lineage ^27^.

Alternatively, LER2 and LER4 have likely fused in the tetrapod ancestor with chicken GGA2 arising from this fusion, and LOC9 and LOC11 where each show the same CLG content than GGA2 are secondarily split of this ancestrally fused chromosome. This may have involved a Robertsonian fission mechanism that effectively splits a metacentric chromosome at the centromere into two acro- or telo-centric products. We also observe several cases in which microchromosomes have been added to macro-chromosomes relatively recently by terminal translocation, such as the addition of a chromosome similar to LER35≡GGA22 to the start of LOC1, or a LER12-like chromosome to the end of GGA4 (a recent translocation not found in other birds such as turkey) ^28^.

The extensive conservation of chromosomal identity and gene order between the little skate and the bamboo shark ^29^, despite over 300my of divergence, indicates that most chondrichthyans share this ancestral condition (**Fig 1b, c**). Interestingly, gene order collinearity across cartilaginous fish is more extensively conserved than within clades of comparable divergence, such as mammals and frogs ^30^. Conversely, gene order is heavily disrupted between chondrichthyans (such as skate or shark) and osteichthyians (gar or chicken in **Fig 2a, b** and **Extended Data Fig. 3**).

The relatively large number of elasmobranch chromosomes (≥40) therefore reflects the ancestral condition among gnathostomes: at the exception of the losses of 2 ancestral segments in the skate lineages, and one secondary fusion in chromosome 1, the skate possessed 37 of the 39 ancestral vertebrate linkage (**Extended Data Table 3**). The evolution of reduced chromosome number in various bony fish lineages is therefore likely due to successive chromosomal fusions.

### Evolution of the gene complement

As in other chondrichthyans, the little skate gene complement has evolved more slowly than that of osteichthyans with respect to the rates of gene losses (**Extended Data Fig. 1**). By reconstructing gene families and applying species-tree aware phylogenetic methods, we found that the retention of ohnologs, the paralogs derived from the vertebrate-specific WGDs, was higher than observed in bony fishes (**Fig. 2c,e**). According to the auto-then-allotetraploidy scenario for jawed vertebrate evolution ^24^ the chromosomes derived from the 2R (the ‘alpha’ and ‘beta’ segments) behave distinctly, with the ‘beta’ segments showing increased loss as well as faster rates of molecular evolution (**Fig. 2c, d,e**).

By querying gene trees based on patterns of duplication and loss, we found 68 cases of asymmetric ohnolog loss, when one ohnologue was differentially retained in varying jawed vertebrate lineages. We also found 19 cases of asymmetric loss retained in chondrichthyans but lost in bony fishes, 17 retained in chondrichthyans and coelacanth, and 24 retained in chondrichthyans and actinopterygians (ray-finned fish) but lost in lobe-finned fish (24 cases) (**Extended Data Table 4**). Some of these retained ancestral ohnologues, including previously characterized genes like *Wnt6b* ^21^, or novel ones such as the chondroitin sulfate proteoglycan 5 (CSPG5) gene shows a distinct expression pattern among stages and organs (**Figure 2f**).

### Conservation of 3D regulatory principles across vertebrate evolution

We investigated features of three-dimensional (3D) chromatin organization in skates using high-throughput chromosome conformation capture (Hi-C) from developing pectoral fin samples. At the level of nuclear organization, we found a “type-II” architecture ^9^ with chromosomes preferentially occupying individual territories within the nucleus (**Extended Data Fig. 4**). This mode of nuclear architecture is consistent with the presence of a complete set of condensin II subunits (*Smc2, Smc4, Caph2, Capg2* and *Capd3*) encoded in the skate genome. At higher resolution, we find that chromosomes are organized into two distinct compartments, analogous to the A/B compartmentalization consistent with their presence in several vertebrate and invertebrate species ^31^. The A compartment consistently displays a higher gene density, more open chromatin regions and higher gene expression levels than the B compartment (**Extended Data Fig. 5**).

At the sub-megabase scale, the skate genome is organized into Topologically Associated Domains (TADs) with a median size of 800 Kb (**Extended Data Fig. 6a, b**). Aggregate plot analyses revealed that skate TADs associate prominently with chromatin loops that can be observed at the upper corner of domains (**Fig. 3a**), thus denoting stable interactions between boundary regions. Assays of chromatin accessibility (ATAC-seq) in both the anterior and posterior region of the skate pectoral fin shed light on the mechanisms of TAD formation. Motif enrichment analysis revealed that binding sites for the architectural factor CTCF are consistently found at TAD boundaries (**Extended Data Fig. 6e**). Importantly, these CTCF binding sites display a clear orientation bias with motifs oriented towards the interior of the TADs, which is consistent with the formation of TADs through loop extrusion (**Fig. 3b, Extended Data Fig. 6e**). A prominent example of skate TAD organization can be observed at the *HoxA* and *HoxD* clusters (**Fig. 3c; Extended Data Fig. 6d**), which display the bipartite TAD configuration that is characteristic of jawed vertebrates ^32^. Importantly, manual microsynteny analysis confirmed that the 3’ and 5’ TADs found at both skate *Hox* loci are homologous to the 3’ and 5’ TADs already described in mammals and teleosts. Such deeply conserved 3D organizations reflect the existence of regulatory constraints that influenced the evolution of TADs across the whole jawed vertebrate clade.

**Fig.3:**
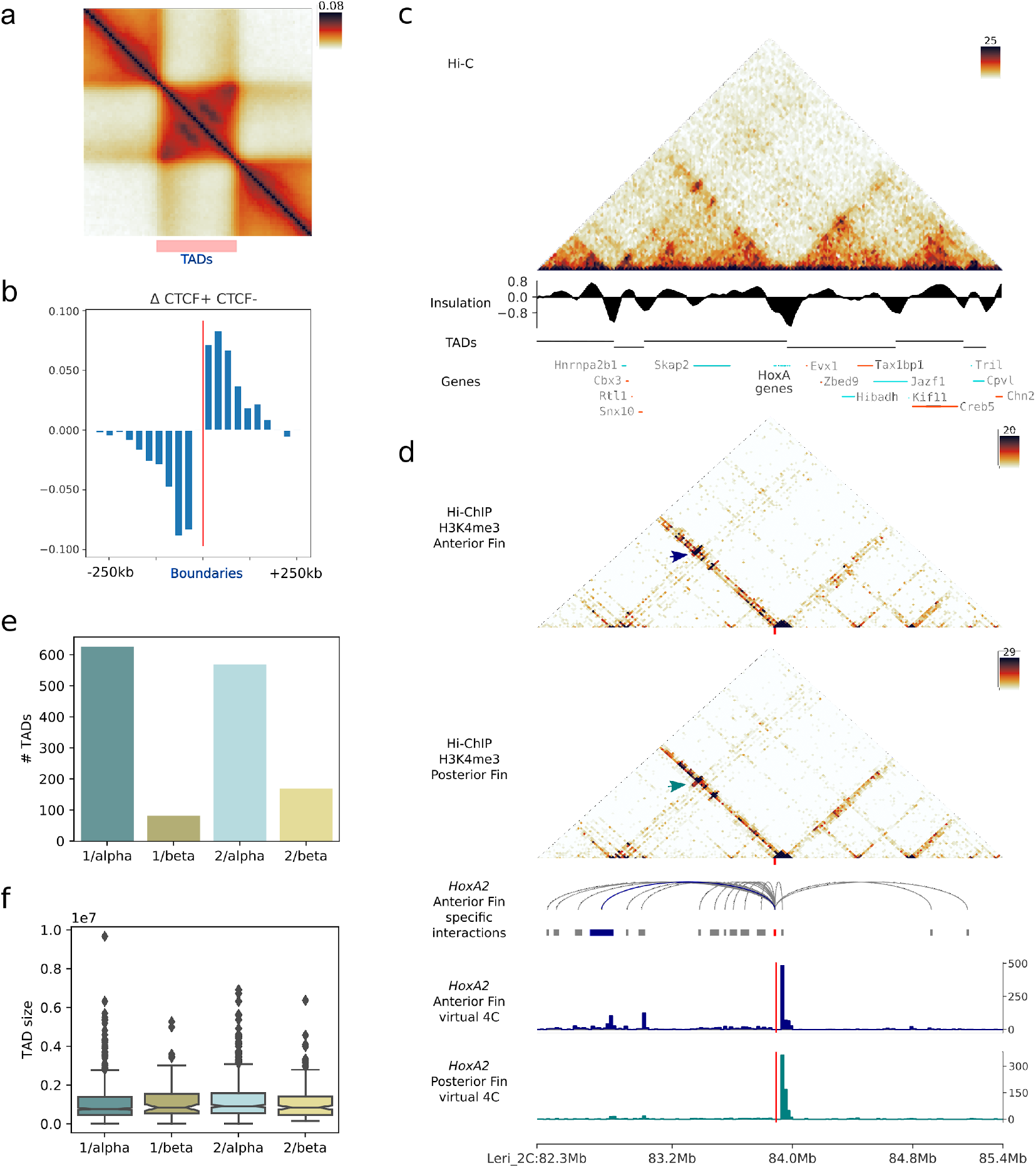
Features of 3D chromatin organization in the little skate. **a**, TAD metaplot displaying focal interactions at the apex of domains. **b**, Orientation bias of CTCF binding site motifs inside ATAC-seq peaks at TAD boundary regions. **c**, Hi-C maps from pectoral fins of the skate *HoxA* locus, denoting the presence of bipartite TAD configuration. **d**, HiChIP H3K4me3 maps from anterior and posterior pectoral fins, displaying tissue-specific interactions (blue line and box in the arachnogram, highlighted with arrows in the HiChIP matrix). Virtual 4C plots obtained from anterior and posterior HiChIPs are shown at the bottom. **e**. Number of TADs detected associated to the different paralogous segments descending from the two rounds of WGD (1 or 2 for the 1R, alpha or beta for the 2R) **f**. TAD sizes observed in the different paralogous segments from **e**.

To further explore enhancer-promoter interactions, we used chromatin-conformation-capture-immunoprecipitation (HiChIP) to associate H3K4me3-rich active promoters with potential regulatory loci in the anterior and posterior region of the developing pectoral fin, where *HoxA* and *HoxD* genes are expressed ^3^. In total we identified 50,601 interactions formed by 7,887 different promoters (median of 6.4 interactions per active promoter). As expected, interactions connecting promoters with distal ATAC-seq peaks (χ^2^ p-value < 10^−138^, **Extended Data Fig. 7a**) and also intra-TAD interactions were enriched (empirical p-value < 10^−4^, **Extended Data Fig. 7b**). Differential analysis revealed similar looping patterns between the anterior and the posterior region of the fin (Pearson correlation >0.96, with 9 and 5 interactions statistically enriched in the anterior and posterior fins respectively, **Extended Data Fig. 7e, f)**.

We found striking differences in promoter interaction between anterior and posterior fins at specific loci including the *HoxA* locus **(Fig. 3d, Extended Data Fig. 7h, i)**. In the anterior fin, the *HoxA2* promoter displays enriched interactions with regulatory regions from the 3’ TAD. Such interaction changes are consistent with the specific expression of *HoxA* genes in the anterior portion of the developing pectoral fin ^3^. Importantly, the presence of interactions between *HoxA* genes and potential regulatory regions in both the 3’ and the 5’ TADs suggests that regulatory constraints maintained the configuration in two TADs of *Hox* loci through jawed vertebrate evolution.

We further explored the relationship between TAD structure and genome evolution arising from regulatory constraint. We found 1,464 microsyntenic pairs of genes (i.e., consecutive orthologs) between skate, mouse and garfish. As expected, such conserved pairs of genes were contained within the same skate TAD more often than other pairs of consecutive genes (98% vs 95%, χ^2^ p-value 3.7 × 10^−13^, **Extended Data Fig. 8a**). Those 1,464 adjacent gene pairs conserved across jawed vertebrates were present in 718 out of the 1,678 TADs identified in the skate genome (around 42%), highlighting that individual TAD structures are constrained but not fully invariant across vast evolutionary timescales (**Extended Data Fig. 8b**). TADs containing deeply conserved microsyntenic pairs are significantly larger and contain more distal ATAC-seq peaks and putative promoter-enhancer interactions, as defined with HiChIP, than non-conserved TADs (**Extended Data Fig. 8c**, Mann-Whitney U p-values of 1.23 × 10^−24^, 3.81 × 10^−36^ and 1.04 × 10^−41^ respectively), suggesting that the cases of deep conservation of individual TADs emerge as a consequence of regulatory constraints (see also *Foxc1* and *Ptch1* loci, **Extended Data Fig. 8d, e**).

Our results suggest that 3D chromatin organization in skates results from the interplay of two mechanisms, compartmentalization driven by transcriptional state and TAD formation mediated by loop extrusion. The similarities in genome organization between skate and bony fishes/tetrapods indicate that the mechanism of TAD formation through loop extrusion was already present in the gnathostome ancestor. Since the appearance of this common ancestor is temporally close to the 2R, we explored the regulatory fate of homologous TADs in relation to this duplication event. We find that although the size of TADs is similar between alpha and beta chromosomes, there are notably fewer TADs in the latter (**Fig. 3e, f**). Regulatory landscapes devised from H3K4me3 HiChIP experiments followed a similar trend compared to TADs (**Extended Data Fig. 7c, d**). These results indicate that a high proportion of TADs disappeared from the early gnathostome genome after the second vertebrate WGD event, while those that are preserved as ohnologues are comparable in size (**Fig. 3f**) and gene content (**Extended Data Fig. 6c**).

### TAD rearrangements identify the PCP pathway as a driver of skate anterior fin expansion

We explored whether TAD-altering genomic rearrangements could have driven skate pectoral fin evolution through gene regulatory changes, as recently reported for other mammalian traits ^12^. To that end, we identified breaks in synteny by aligning six jawed vertebrate genomes, including two skates, two sharks, a chimera, and gar fish; **Methods, Fig. 4a**) As expected, the number of (micro)syntenic changes between species detected increases with phylogenetic distance (**Fig. 4a, Extended Data Fig. 9a**), ranging from the 18 breaks in *Leucoraja erinacea* that occurred after the split of the two skate lineages to the 1,801 differences identified between cartilaginous and bony fishes, which correspond to a rate of ∼2 breaks per million years.

**Fig.4:**
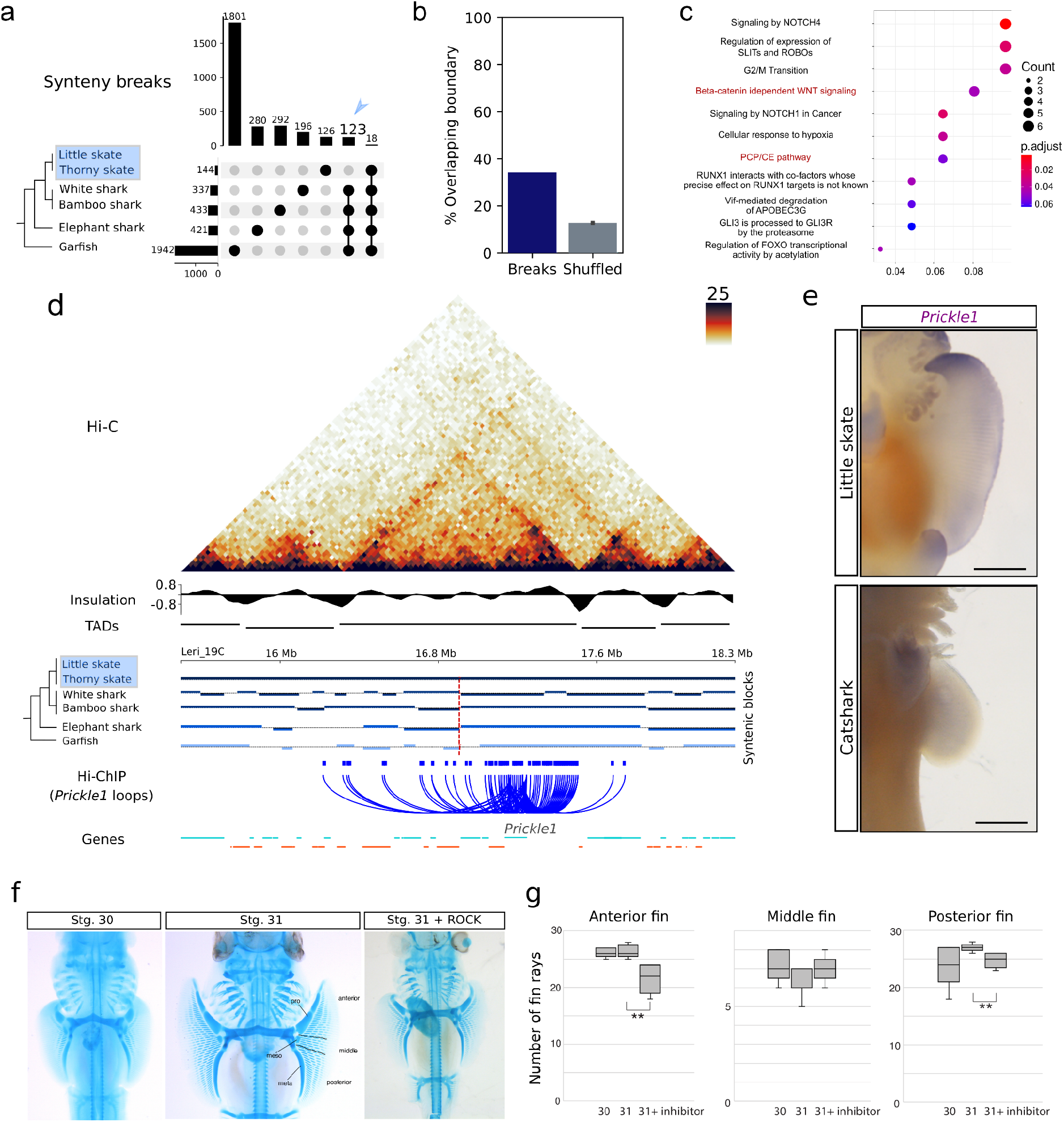
Skate-specific genomic rearrangements and the PCP pathway. **a**, Synteny breaks detected in skates and related vertebrates. Blue arrow indicates the 123 genomic rearrangements that are specific for skates. **b**, Percentage of synteny breaks that appear at TAD boundaries (dark blue) and expected percentage for shuffled boundaries (grey). **c**, Reactome signaling pathway analysis on genes contained in rearranged TADs. **d**, Hi-C map from pectoral fins of the *Prickle1* locus. Synteny blocks are indicated, together with insulation scores, TAD predictions and chromatin loops detected in H3K4me3 HiChIP datasets. **e**, Whole-mount *in situ* hybridization for *Prickle1* in skates (*Leucoraja erinacea*, stage 30) and catshark (*Scyliorhinus retifer*, stage 30). Note that anterior expression is specific to skates. **f**, Cartilage staining of embryos with/without a rho-kinase (ROCK) inhibitor. Compared to negative control embryos at stage 30 and 31, the number of fin rays decreased in the embryos with the ROCK-inhibitor. Note the reduction of fin rays on the anterior region of the pectoral fin is severer than that on the posterior region. Photos of all replicates are shown in Extended Data 13-15. The pectoral fin was divided into three domains from the anterior to the posterior (see Methods). pro; propterygium, meso; metapterygium, meta; metapterygium.

Since anterior expansion of the pectoral fin is a defining characteristic of skates, we focused on the 123 synteny breaks shared by the little and thorny skate genomes relative to other jawed vertebrates. We found an enrichment of synteny breaks near TAD boundaries: 42 breaks occurred within 50 kb of a TAD boundary, compared with 15 expected under a random break model (empirical p-value < 1×10^−4^; **Fig. 4b**). This nearly 3-fold enrichment supports the hypothesis that genome rearrangements that interrupt TADs are evolutionarily disfavored because they typically cause deleterious enhancer-promoter rewiring ^12^.

Conversely, we hypothesized that the 81 breaks that do interrupt TADs could be enriched for enhancer-promoter rewiring associated with skate-specific changes in gene regulation. TADs interrupted by breaks include a total of 2,041 genes. We identified a restricted set of 180 genes likely associated with pectoral fin expansion by intersecting this list with genes displaying interactions across the synteny breaks as detected by H3K4me3 HiChIP on anterior fins. Signaling pathway enrichment analysis revealed the enrichment of components of the Wnt/PCP pathway (**Fig. 4c, Extended Data Fig. 9b, c**) including the important regulator *Prickle1* (**Fig. 4d**). Other potentially relevant genes including the known *Hox* gene activator *Psip1* were also identified ^33^ **(Extended Data Fig. 10**).

To test our hypothesis that changes in chromatin architecture drove changes in gene expression patterns we performed comparative *in situ* hybridizations of *Prickle1* in paired fins of the little skate and the chain catshark (*Scyliorhinus retifer*) at the equivalent stage 30 (**Fig. 4e**). Consistent with the synteny break data, we detected a distinct expression pattern between the pectoral fins of skate and shark. Specifically, *Prickle1* displayed higher expression in the anterior pectoral fin of the little skate compared to a weak expression without any spatial enrichment in shark fins (RT-PCR result in **Extended Data Fig. 11**). Similarly, we also found differential expression for *Psip1*, which could suggest a potential involvement of *Hox*-related pathways in the skate fin phenotype (**Extended Data Fig. 10c, d**).

Given the specific pattern of *Prickle1* expression, we further explored the function of the PCP pathway in the anterior fin extension through cell shape analyses, which revealed that anterior mesenchymal cells are more oval compared to the cells in the central and posterior fin (**Extended Data Fig. 12**). We tested the functional role of the PCP pathway by analyzing pectoral fin formation in skate embryos treated with a Rho-kinase (ROCK) inhibitor from stage 29 to 31. These analyses revealed that the overall number of fin rays was reduced in the ROCK-inhibited embryos compared to the control embryos, with greater losses in the anterior part of the fin compared to the posterior (**Fig. 4f, Extended Data Fig. 13-15**). Although there was significant variation within each stage and treatment (**Extended Data Fig. 13 and 15**), geometric morphometric analyses suggest that, in contrast to control embryos where the pectoral fin extends anteriorly towards the eye by stage 31, ROCK-inhibitor treated embryos showed a less pronounced anterior expansion of the pectoral fin (**Extended Data Fig. 16**). Taken together, these findings suggest that TAD rearrangements contributed to recruit and repurpose genes and pathways responsible for the evolution of the exceptionally unique pectoral fin morphology in a common ancestor of batoids.

### HOX-mediated *Gli3* repression as a mechanism for the formation of a secondary organizer domain in skate pectoral fins

To further explore transcriptional drivers of the derived pectoral fin morphology, we compared RNA-seq datasets from the pectoral fin against the pelvic fin, which exhibits a characteristic tetrapod gene expression pattern ^3^. We identified 193 and 117 genes preferentially expressed in pectoral and pelvic fins, respectively (**Extended Data Table 5**), including several TFs and components of different signaling pathways. To identify potential changes in the appendage gene regulatory network (GRN), we compared our list of differentially expressed genes in skate fins against an analogous gene list resulting from comparing mouse fore- and hindlimb RNA-seq datasets (**Fig. 5a; Extended Data Fig. 17**). Key genes in determining anterior and posterior paired appendages, such as *Tbx5* and *Tbx4*, display a conserved expression pattern, suggesting a conserved patterning function across jawed vertebrates ^34^. However, several genes displayed clear differences between skates and mice (**Extended Data Fig. 17**), suggesting that GRN changes may contribute to the evolution of appendage morphology. A striking example is the expression of *Pitx1*, a master regulator of vertebrate pelvic appendage specification ^35,36^, which was expressed also in both pectoral fins (**Extended Data Fig. 17, 18**). These findings suggest that altered regulation of appendage-related factors may be a major contributor to paired fin expansion in skates.

**Fig. 5:**
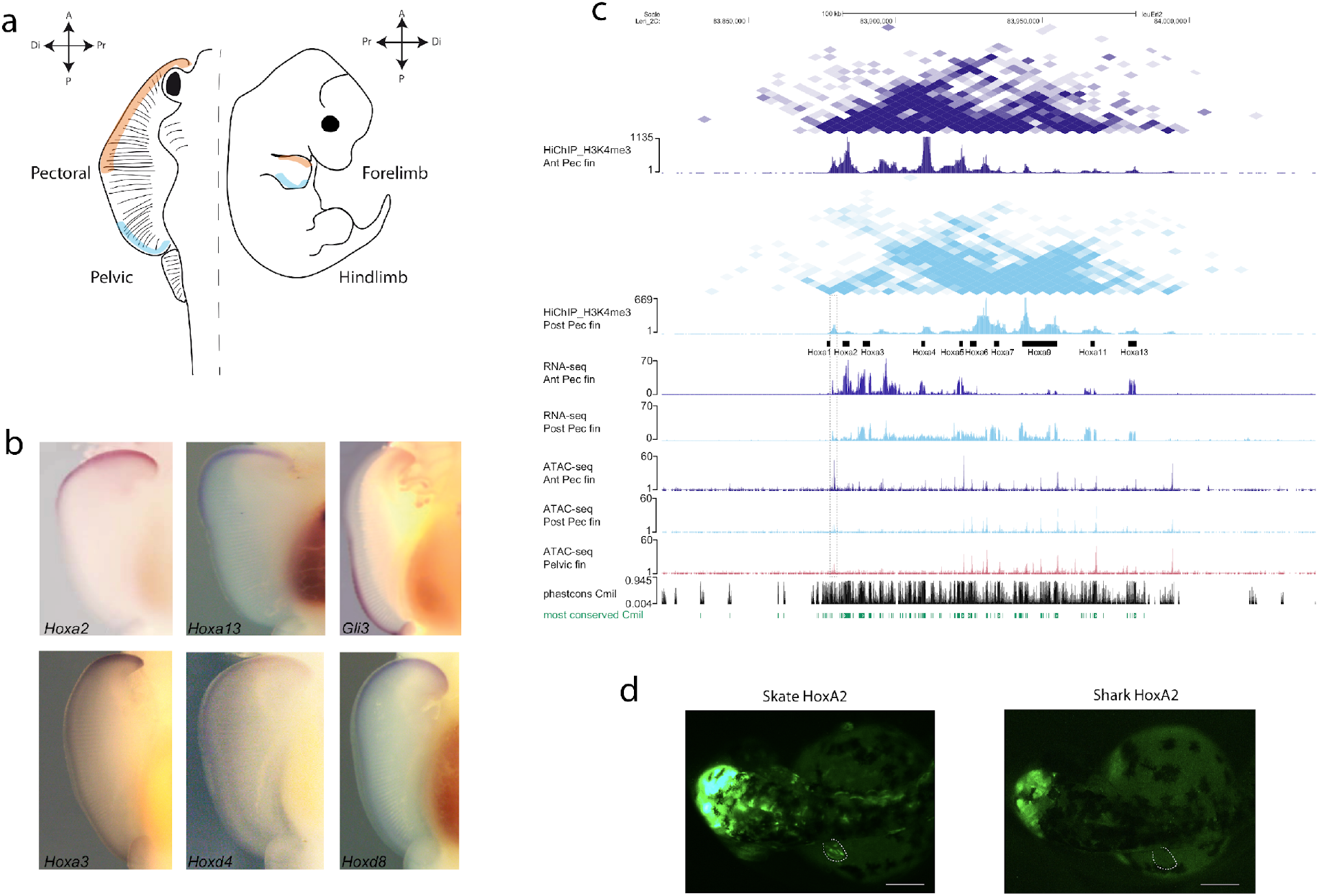
Functional experiments in skate fins samples. **a**. Diagram of the skate fins used to generate our data series, showing homologous structures in pectoral fin and mouse forelimb. A stands for anterior, P for posterior, Di for distal and Pr for proximal regions of the fin/limb. **b**. *In situ* hybridization technique reveals the opposite expression pattern of many *Hox* genes and the *Gli3* gene in the pectoral fin. The images of *Hoxa2* and *Gli3* are reused from ^3^. **c**. UCSC Genome Browser view showing HiChIP, RNA-seq and ATAC-seq data around *Hoxa* cluster in skate. The anterior-specific open chromatin region between *Hoxa1* and *Hoxa2* genes is marked with a dotted rectangle (also see the text). **d**. GFP expression driven by the anterior-specific open chromatin region between *Hoxa1* and *Hoxa2* genes from skate and shark in transgenic assays in zebrafish. Note that only the skate enhancer drives expression on the pectoral fin.

Importantly, *Hox* genes also show distinctive differences between pectoral and pelvic fins (**Extended Data Table 5)** and between mouse and skate paired appendages (**Extended Data Fig. 17**). In particular, several 3’ *HoxA* and *HoxD* genes are preferentially expressed in the anterior pectoral fin, while 5’ *HoxA* and *HoxD* genes are located in the posterior pectoral domains, consistent with previous findings in two different skate species ^3,37^. The anterior expression of *HoxA* and *HoxD* genes is responsible for the development of a secondary apical ectodermal ridge-like organizer, likely involved in the anterior overgrowth of the batoid pectoral fins ^3,37^. The formation of the secondary (AER) is associated with changes in the spatial expression pattern of *Gli3*, a key regulator of Hedgehog signaling involved in appendage patterning ^38,39^. Specifically, *Gli3* is expressed in the posterior domain of the pectoral fins, while in pelvic fins the expression is predominantly anterior, as reported for several vertebrate species. (**Fig. 5b**, ^3^. Recently, it has been shown that *HoxA13* and *HoxD13* genes are required to downregulate *Gli3* expression for proper thumb formation ^40^ in the mouse limb. In addition, Hox13 proteins bind to and repress *Gli3* limb enhancers, and compound Hox13 mutants cause anterior extension of *Gli3* expression. This opens the possibility that the expression of several *Hox* genes in the anterior region of skate pectoral fin may be causing *Gli3* repression that, consequently, contributes to the striking expansion of skate fins. Notably, Hoxa2 can bind to several ehnhancers within the Gli3 locus (as shown by ChIP-seq data ^41^ see **Extended Data Fig. 19**), and is one of the most strongly expressed genes in anterior pectoral fins (**Extended Data Fig. 17** ^3^). Altogether, these results indicate that *Gli3* downregulation mediated by Hox proteins may be one of the major determinants for the skate pectoral phenotype.

To further test this hypothesis, we employed a *hoxd13a-GR* overexpression construct in zebrafish, which becomes active upon addition of dexamethasone ^42^. Using this approach, we had previously demonstrated that the induced overexpression of *hoxd13a* in zebrafish caused increased fin proliferation, distal expansion of chondrogenic tissue, and fin fold reduction. Upon dexamethasone treatment, we further observed that 35% of the injected zebrafish embryos showed reduction of *gli3* expression in the fins (**Extended Data Fig. 20**). In concordance with this observation, a recently generated *Gli3* loss-of-function mutant in medaka fish shows multiple radials and rays in a pattern largely similar to the polydactyly of mouse Gli3 mutants, but also to pectoral skate fins ^43^. Altogether, these findings support *gli3* downregulation, mediated by HOX repression, as a potential mechanism underlying the striking pectoral skate fin shape.

### A skate fin-specific enhancer is linked to the redeployment of *HoxA* gene expression in anterior pectoral fins

Since many of the antero-posterior expression differences found in other vertebrate appendages are not observed in skates, we hypothesized that they may result from changes in regulation. To identify *cis*-regulatory elements (CREs), we performed ATAC-seq in anterior and posterior pectoral fins, as well as in whole pelvic fins. We focused on CREs associated with differentially expressed genes, based on HiChIP datasets. This approach identified hundreds of differentially accessible ATAC peaks, many of those clustered around a small subset of genes that are critical for appendage patterning such as *Tbx5, Tbx4, Pitx1* and multiple *Hox* genes (**Extended Data Table 6**). Of note, this combined analysis revealed that *Pitx1* displays a largely similar regulatory landscape in pectoral and pelvic fins (**Extended Data 18, 20**), which contrasts with the tissue-specific regulatory changes observed in mouse ^36^. Overall, these datasets constitute a powerful resource for identifying potential CREs specifically controlling the expression of genes in pectoral and pelvic fins.

To further determine the regulatory mechanisms behind the anterior expression of Hox genes in skate pectoral fin, we integrated our anterior and posterior pectoral fin ATAC-seq data with the previously published RNA-seq datasets generated in these tissues ^3^. With this approach, we only found a few differentially accessible enhancers associated with differentially expressed genes, but all of them were relevant for anterior-posterior patterning such as *HoxA2, Pax9, Tbx2* and *Alx4* in the anterior pectoral fin, as well as *Chordin, HoxA9, HoxD10, HoxD11, HoxD12* and *Grem1* in the posterior region (**Extended Data Table 5**). Intriguingly, we identified a region closely located to *Hoxa2* that is distinctively more accessible in the anterior region of pectoral fins than in the posterior region or in pelvic fins (**Fig. 5c**). Functional analysis using zebrafish transgenic assays showed that this open chromatin region indeed shows enhancer activity and promotes expression in the anterior region of the zebrafish developing pectoral fins (**Fig. 5d**), compatible with the expression of several *HoxA* genes in the anterior territory of skate pectoral fins (**Fig. 5b**). Sequence conservation analysis revealed that this enhancer is conserved in chondrichthyes but is not found in other bony fishes or vertebrate species (**Extended Data Fig. 23**). Nevertheless, the orthologous region in catshark does not promote transgene expression in functional analyses in zebrafish (**Fig. 5d**), suggesting that, although this enhancer is shared by different chondrichthyan species, only the batoid sequence is functionally active during early development. Since this potential enhancer lies very close to the *HoxA2* promoter, we wondered whether it might be specific for this gene, or it is rather shared with others. Using the H3K4me4 HiChIP data, we observed that this enhancer forms robust interactions with most genes of the cluster in the anterior pectoral fins (**Fig. 5c**), including *Hoxa13* that is located in the 5’ adjacent TAD (**Fig. 3c and 5c)** and expressed in the anterior pectoral fin (**Extended Data Fig. 17**). Overall, these results demonstrate the existence of skate-specific CREs that can be directly linked to the formation of a secondary apical ectodermal ridge-like domain in the anterior margin of the skate pectoral fin.

Finally, DNA methylation profiling (**Extended Data Fig. 22a**) of embryonic and adult pectoral and pelvic fins revealed that differentially accessible ATAC peaks are hypomethylated in developing pectoral and pelvic fins and remain hypomethylated in adult fins (**Extended Data Fig. 22b, c**). This finding suggests that these cis-regulatory elements (CREs) might retain epigenetic memory of developmental activity in adult fins, similar to mouse ‘vestigial’ enhancers and CpG-rich enhancers that participate in body plan formation in zebrafish ^44,45^. Notably, these skate CREs exhibited elevated CpG density in agreement with their hypomethylated state (**Extended Data Fig. 22b, c**). It will be interesting to decipher in future studies whether the persistent hypomethylated state of CREs in adult fins enables recovery of embryonic epigenetic memory that could potentially drive CRE reactivation during fin regeneration or other biological processes.

## DISCUSSION

Through a suite of genomic analyses and functional validations, we uncover fundamental principles of genome regulation in the skate lineage, and provide a molecular basis for the formation of wing-like fins in batoids ^1^. The convenient position of skates in the vertebrate evolutionary tree, as well as their slower rate of genome evolution provided important insights on karyotype stabilization after two rounds of whole genome duplication [2R]. We find that gene loss and karyotype evolution dynamics have occurred at a different pace across jawed vertebrate lineages. While most previous studies have explored the consequences of genome duplications in tetrapods and teleosts, the elephant shark genome exhibited a slower rate of evolution and reduced gene loss compared to tetrapods ^26,46^. Here we extend this observation to the scale of chromosomes, and show that the skate genome exhibits extensive retention of ancestral gnathostome chromosomes: the fewer chromosomes of chicken and spotted gar were formed by fusion of these ancestral units. This process seems accompanied by considerable gene order rearrangement between cartilaginous fishes (chondrichthyans) and bony fishes (osteichthyans), which contrasts with the extensive conservation of TAD gene contents at long evolutionary distances. This conservation in the absence of a globally colinear gene order emphasizes the impact of regulatory constraints in maintaining some of these gene groupings.

Strikingly, we also observed the complete disappearance of entire TAD units in the paralogous regions prone to gene loss after the second WGD (the ‘beta’ segments) for which we detect a lower number of TADs, but with the same average size and gene number. While asymmetric paralog loss following WGDs is considered a key factor in the emergence of novel gene regulations ^5^, the loss of TADs in ‘beta’ regions indicates that entire paralogous regulatory units can be lost after WGDs and stresses the importance of regulatory constraints in shaping genome organization. It remains to be demonstrated if the regulatory potential of the missing TADs is incorporated into other regulatory landscapes, thus contributing to additional gene pleiotropy. We also find that the skate genome is functionally constrained by mechanisms of 3D gene regulation that are analogous to those described in bony fishes and tetrapods. These include the presence of a CTCF-orientation code and associated loop extrusion ^14^. Importantly, our findings suggest that these structural mechanisms emerged early in vertebrate evolution, likely influencing the appearance of their phenotypical novelties. These mechanisms induce further constraints as most skate-specific chromosome rearrangements occur at TAD boundaries, resulting in limited effects on gene regulation, as also reported in mammals ^47^. Nevertheless, TAD-disrupting rearrangements are also detectable, affecting several genes involved in the PCP pathway, an ancient developmental pathway whose function can be traced back to Porifera ^48^. The PCP pathway is essential for cell orientation and patterning, as demonstrated by functional experiments resulting in altered limb morphogenesis, and its disruption is the molecular basis of certain human congenital malformations. Thus, the anterior-specific modulation of the PCP pathway, through a tissue-specific redeployment of a main pathway effector (*Prickle1*), provides a compelling example of how existing gene networks can be reutilized to evolve new functions during animal development through genomic regulatory rearrangements.

These results are complemented with our findings on the regulation of 3’*Hox* genes, as well as their activator *Psip1*. These genes are expressed in the anterior pectoral fins in contrast to their posterior expression observed in most vertebrate appendages or even in skate pelvic fins. Based on our *hoxd13a* overexpression experiments (**Extended Data Fig. 17**), we suggest that the increased level of *Hox* genes expression, specifically in the anterior pectoral fin, together with other regulatory changes, contributes to the downregulation of *Gli3* in this territory which may be one of the major drivers of the striking morphology of skate pectoral fins. This result illustrates the remarkable plasticity of the *Shh/Gli3/Ptch1* pathway in contributing to the evolution of vertebrate appendage morphology ^43,49–51^. In addition, our epigenomic and functional analysis using reporter assays in zebrafish identified a *HoxA* fin enhancer specific to the skate genome, which provides an additional regulatory basis for such patterning variations on signaling pathways. Overall, our study on the evolution of the batoid fin exemplifies how variations on regulatory elements and in 3D chromatin organization can act as major evolutionary forces for species adaptation.

## METHODS

### Genomic DNA extraction and library construction

Skate DNA was isolated using extensive proteinase K digestion and Phenol-Chloroform extraction from the muscle of a single Leucoraja erinacea specimen. For genome assembly, we generated both accurate short-reads and noisy long-reads. A contiguous long read (CLR) library for Pacbio sequencing was prepared and sequenced at the Vincent J. Coates Genomics Sequencing Laboratory at UC Berkeley. A total of 32 cells were sequenced on the Pacbio Sequel instrument using the V7 chemistry and yielded a total 10.2M of pacbio reads totalling 163 Gb with a median size of 10.9 kb and a read N50 of 29kb.

A paired-end illumina library with a 600 bp insert was also sequenced for 2×250 bp in rapid run mode on a HiSeq 2500 instrument at BGI (Guangdong, China) yielding 641M reads and 160.3Gb of sequence.

### Genome assembly

Genome size was estimated by analyzing a k-mer spectrum with a mer size of 31. By fitting a multimodal distribution using Genomescope 2.0, and estimated a genome size of 2.13Gb (as well as an heterozygosity of 1.56%) ^52^. To take advantage of both short and long reads, we opted for a hybrid assembly strategy. First, we generated *de brujin* graph contigs using megahit (v1.1.1) using a multi-kmer approach (31, 51, 71, 91 and 111-mers) and filtering out k-mers with a multiplicity lower than 5 (--min-count 5). We obtained 2,750,419 contigs with a N50 of 1,129bp representing a total of 2.23Gb. We then used these contigs to prime the alignment and assembly of the pacbio reads using dbg2olc (c. 10037fa) ^53^ using a k-mer of 17 (k 17), a threshold on k-mer coverage of 3 (*KmerCovTh 3*), a minimal overlap of 30 (*MinOverlap 30*) and an adaptive threshold of 0.01 (*AdaptiveTh 0*.*01*) and removing chimeric reads from the dataset (RemoveChimera 1). This assembler generated an uncorrected backbone of overlapping reads with a N50 of 4.96Mb and a total size of 2.25Gb. To correct sequencing errors, we subjected this sequence file to two successive rounds of consensus by aligning pacbio reads with minimap2 (v. 2.12, *map-pb* setting) ^54^ and Racon (v. 1.3.1) with default parameters followed by one final round of consensus using the illumina reads. We evaluated the progress of the polishing process with the BUSCO tool (v. 3.0.2) that seeks widely represented single-copy gene families in the assembly ^55^. Our final polished assembly contained 95.1% of vertebrate BUSCO genes (**Extended Data Table 1**). To exclude residual haploid contigs from the assembly, we aligned illumina reads once more using bwa and computed a distribution of coverage that showed some residual positions at half coverage (31x). We used purge_haplotigs ^56^ by defining a coverage threshold between haploid and diploid contigs at 40x (and a minimum of 10x and maximum of 100x). The filtered assembly has a size of 2.19Gb, a N50 of 5.35Mb and 2595 contigs in total and the same BUSCO statistics as the unfiltered one (**Extended Data Table 1**).

This assembly was then scaffolded using chromatin-contact evidence obtained from H-iC sequencing of *L. erinacea* fins (see below) at Dovetail genomics (Scotts Valley) with the HiRise pipeline ^57^. The accuracy of the resulting scaffolded assembly was verified and proofread by carefully inspecting the contact map in Juicebox ^58^ and HiGlass browser ^59^. This assembly comprises 50 scaffolds larger than 1Mb that represent 92% of the assembly size and 39 scaffolds larger than 10Mb that show mostly internal contacts. Despite no karyotyping evidence is directly available for *L. ericinea*, closely related species show a haploid number of 49 chromosomes, which is consistent with the observed number of chromosomes ^15^.

As the final assembly size was smaller than the experimentally assessed genome size of 3.5Gb, we perform gap-closing on the final assembly using PBjelly ^60^ that proceeds through alignment of the pacbio reads on each gap border and local reassembly. The impact on the assembly statistics was marginal, but we used this assembly as our final one (Table S1).

### Annotation

RNA-seq reads of strand-specific libraries from five bulk embryonic stages were aligned to the genome using STAR (v. 2.5.2b) ^61^ and each library assembled independently using stringtie (v. 1.3.3) ^62^. Stringtie assemblies were then merged using TACO (v. 0.7.3) ^63^. RNA-seq reads were also assembled *de novo* using Trinity (v. 2.8.4) ^64^. Finally, the iso-seq protocol was applied to generate full-length transcripts using Pacbio long-reads. Both Trinity assembled transcripts and iso-seq transcripts were aligned to the genome using GMAP (v. 2018-07-04) ^65^. Then, both TACO assembled transcripts and aligned *de novo* transcripts were leveraged using Mikado (v. 1.2.1) ^66^ to generate one consensus reference transcriptome, while predicting coding loci using Transdecoder (v. 5.5.0). Using selected transcripts (2 introns or more, complete CDS, single hit against swissprot), we build an Augustus HMM profile for *ab initio* gene prediction ^67^. We predicted skate genes using this profile and hints derived from (1) the mikado transcript assembly (exon hints), (2) intron hits obtained using bam2hints on a merged bam alignment of the RNA-seq data after filtering spurious junctions with portcullis ^68^ and (3) an alignment of human protein using exonerate ^69^.

A repeat library was constructed using Repeatmodeler and repeats were masked in the genome using Repeatmasker (v4.0.7). We filtered out gene models that overlap massively with mobile elements and obtained 30,489 genes models. For these genes, isoforms and UTRs were added by two rounds of reconciliation with an assembled transcriptome using PASA ^70^. Our set of coding genes includes 5,800 PFAM domains, a similar value than other well-annotated vertebrate genomes. To further examine the validity of gene models, we assessed (i) whether their coding sequence showed similarity to that of another species using gene family reconstruction (see below) (ii) whether they possessed an annotated PFAM domain and (iii) whether they are expressed above 2 FPKMs in at least one RNA-seq dataset. These criteria reduced the number of bona fide coding genes to 26,715.

### Gene family, synteny and phylogenetic analyses

We performed gene family reconstruction using OMA ^71^ between selected vertebrate species to identify single-copy orthologues. These orthologues were used to infer gene phylogeny after processing as described in ^72^: HMM profiles were built for each orthologous gene family and searched against translated transcriptomes using the HMMer tool (v3.1b2) ^73^. Alignments derived from each orthologue were aligned using MAFFT (v7.3) ^74^, trimmed for misaligned region using BMGE (v1.12) ^75^ and assembled in a supermatrix. Phylogeny was estimated using IQTREE (v2.1.1) assuming a C60+R model and divergence times estimated using Phylobayes (v4.1e) ^76^ assuming a CAT+GTR substitution, and a CIR clock model, soft constraints and a birth-death prior on divergence time. Calibrations were derived from ^19,77^.

We identified conserved segments across vertebrates, by counting single-copy copy genes derived from OMA clustering sharing the same set of chromosomal locations in selected species, to identify putative ancestral vertebrate units. We examined conserved syntenic orthology by identifying sets of genes shared by pairs of chromosomes in distinct species using reciprocal best hits computed using Mmseq2 ^78^. We performed a fisher test to detect pairs of chromosomes showing significant enrichment, and assigned ALG based on comparison with amphioxus and sea scallop. We computed gene family composition and analysed patterns of gene loss and duplications using reconstructed gene trees derived from gene families established with Broccoli ^79^ and subjected to species-tree aware gene tree inference using Generax ^80^.

### Hi-C

Hi-C protocol was carried out as described elsewhere with almost no modifications ^81^. Two biological replicates of *L*.*erinacea* Stg.30 pectoral fin buds were fixed in a final concentration of 1%PFA for 10’ at room temperature. Fixation was stopped by placing the samples on ice and by adding 1M glycine up to a concentration of 0.125M. The quenched PFA solution was then removed and the tissue was resuspended in ice-cold Hi-C Lysis Buffer (10mM pH=8 Tris-HCl, 10mM NaCl, 0.2% NP-40 and 1x Roche Complete protease inhibitor). The lysis was helped with a Dounce Homogenizer Pestle A on ice (series of 10 strokes in 10’ intervals). Nuclei were then pelleted by centrifugation for 5’, 2500 rcf at 4ºC, washed twice with 500 μL of 1x NEBuffer3.1 and finally resuspended with the same buffer to final volume of 374 μl. 38 μl of 1% SDS was then added and the sample incubated 10’ at 65ºC. SDS was then quenched with 43 μl of 10% Triton X-100. Chromatin was then digested by adding 12 μl of 10x NEBuffer3.1 and 8 μl of 5 U/μl DpnII (NEB, R0543) followed by a 2h incubation at 37ºC in a ThermoMixer with shaking (900 rpm, 20” pulses every 3’). DpnII was then heat inactivated at 65ºC during 20’ with no shaking. Chromatin sticky ends where then filled-in and marked with biotin by adding 60 μl of Fill-in Master Mix (6 μl of 10x NEBuffer3.1, 1.5 μl of 10mM mix of dCTP, dGTP and dTTP, 37.5 μl of 0.4 mM biotin-dATP (Thermo Fisher, 19524016) and 10 μl of 5 U/μl Klenow (NEB M0210)) and incubating 1h at 37ºC with rotation. Filled-in chromatin was then ligated by adding 970 μl of Ligation Master Mix (150 μl of 10x NEB T4 DNA ligase buffer with ATP (NEB, B0202), 125 μl of 10% Triton X-100, 15 μl of 10 mg/ml BSA and 10 μl of 400 U/μl of T4 DNA ligase (NEB, M0202)) and incubated 4h at 16ºC with mixing (800 rpm, 30” pulses every 4’). Ligated chromatin was then reverse crosslinked adding 50 μl of 10 mg/mL proteinase K and incubating the sample at 65ºC for 2h. Decrosslinking was completed by adding 50 μl extra of proteinase K and incubating overnight at 65ºC. DNA from the reverse crosslinked chromatin was purified using phenol:chloroform extraction and ethanol precipitation. Pelleted DNA was resuspended in 1 ml of TLE and washed twice by dyalisis with 8 ml of TLE using AMICON Ultra Centrifuge Filter Unit 15mL 30K (Millipore UFC903024, around 500 μl of sample was recovered from last centrifugation). Then 1 μl of 10 mg/ml RNAse was added and the sample incubated 30’ at 37”. After RNA degradation, DNA was quantified using Qubit and biotin was removed from unligated ends. Biotin removal was performed in a final volume of 50 μl with 5 μg of the quantified DNA, 5 μl of 10x NEBuffer2.1, 0.125 μl of 10 mM dATP, 0.125 μl of 1 mM dGTP, 5 μl of 3000 U/ml T4 DNA polymerase (NEB M0203L). The sample was incubated in a thermocycler 4h at 20ºC followed by 20’ at 75ºC and then dialysed again using Amicon Ultra-0.5 Centrifugal Filter Unit (Millipore UFC5030) and milliQ water. 130 μl were recovered from the dyalysis columns and used for DNA sonication in a M220 Covaris Sonicator (Peak power: 50, Duty factor: 20%, Cycles/Burst: 200, Duration 3’). After sonication, DNA was size selected with AMPure XP beads (Agencourt, A63881). Briefly, in a first selection, 0.8x of beads were used and the supernatant was recovered. In the second selection, 1.1x of beads were used and the bead fraction was recovered. Size-selected DNA was resuspended in 50 μl of TLE and then subjected to end repair. End repair was performed by adding 20 μl of the End Repair Mix (7 μl of 10x NEB ligation buffer, 1.75 μl of 10 mM dNTP mix, 2.5 μl of T4 DNA polymerase (3 U/μl NEB M0203), 2.5 μl of T4 PNK (10 U/μl, NEB M0201) and 0.5 μl of Klenow DNA polymerase (5 U/μl NEB M0210)) and incubating in a thermocycler with the following program: 15ºC 15’ / 25 ºC 15’ / 75 ºC 20’. Biotinylated ligation ends were then pulled down using 10 μl of Dynabeads© MyOne™ Streptavidin C1 (Invitrogen, 650.01) per μg of DNA. The beads were washed twice with Tween Wash Buffer (85mM Tris-HCl pH=8, 0.5 mM EDTA, 1M NaCl, 0.05% Tween) before being resuspended in 400 μl of 2x Beads Binding Buffer (10 mM Tris-HCl pH=8, 1 mM EDTA, 50 mM NaCl) and incubated 15’ with rotation with 400 μl of the end repaired sample (70 μl of end repair reaction plus 330 μl of TLE). Beads were then washed once with 400 μl of 1x Beads Binding Buffer and once with 100 μl TLE before being finally resuspended in a final volume of 41 μl. A-Tailing was then performed in a total volume of 50 μl by adding 5 μl of 10x NEBuffer2.1, 1 μl of 10mM dATP and 3 μl of 5 U/μl Klenow fragment 3’ -> 5’ exo- (NEB M0212) in the thermocycler with the following program: 37ºC 30’ / 65ºC 20’. A-tailed sample was then washed with 400 μl of 1X T4 ligase buffer and resuspended in 40 μl of the same buffer to prepare it for the adaptor ligation, that was performed adding 1 μl of 10x T4 ligation buffer, 4 μl of T4 DNA ligase and 5 μl of 15 μM Illumina paired-end pre-annealed adapters. The reaction was incubated 2h RT and then the beads were washed twice with 1x NEBuffer2.1. Beads were resuspended in 50 μl of the final library PCR reaction for library generation (25 μl of NEBNext® High-Fidelity 2x PCR Mix, 0.5 μl of PE1 primer 25 μM and 0.5 μl of PE2 primer 25 μM plus milliQ water). The PCR was carried out in a thermocycler with the following program: 72ºC 5’ / 98ºC 30” / 7-10 cycles of 98ºC 10” - 63ºC 30” - 72ºC 30” / 72ºC 5’. Pilot PCRs were used to determine the number of cycles. Final single-sided AMPure XP beads purification was performed to eliminate primer-dimers (1.1x proportion). Final libraries were sent for paired-end sequencing.

### Hi-C analysis

Hi-C paired-end reads were mapped to the skate genome using BWA ^82^. Then, ligation events (Hi-C pairs) were detected and sorted, and PCR duplicates were removed, using the pairtools package (https://github.com/mirnylab/pairtools). Unligated and self-ligated events (dangling and extra-dangling ends, respectively) were filtered out by removing contacts mapping to the same or adjacent restriction fragments. The resulting filtered pairs file was converted to a tsv file that was used as input for Juicer Tools 1.13.02 Pre, which generated multiresolution hic files ^83^. These analyses were performed using already published custom scripts (https://gitlab.com/rdacemel/hic_ctcf-null): the hic_pipe.py script was first used to generate tsv files with the filtered pairs, and the filt2hic.sh script was then used to generate Juicer hic files. Visualization of normalized Hi-C matrices and other values described below, such as insulation scores, TAD boundaries, aggregate TAD, Pearson’s correlation matrices and eigenvectors, were calculated and visualized using FAN-C ^84^ and custom scripts present in the git repository https://github.com/skategenome. The observed-expected interchromosomal matrix (**Fig. 1d**) was calculated counting interchromosomal normalized interactions in the 1Mb KR normalized matrix (with the two replicates merged). Expected matrix was calculated as if interchromosomal interactions between two given chromosomes were proportional to the total number of interchromosomal interactions of these two chromosomes. A/B compartments were first called in each of the replicates separately using the first eigenvector of the 500kb KR normalized matrix. Eigenvector correlation was high (r=0.91, **Extended Data Fig. 7b**) and the replicates were then merged. The first eigenvector was calculated again and oriented according to open chromatin using the amount of ATAC-seq signal in the anterior pectoral fin sample. ATAC-seq, %GC, gene models and RNA-seq signal overlaps with compartments were calculated using *bedtools intersect* ^*85*^. Compartment calling and the different overlaps are available in **Extended Data Table 8**. The saddle plot was calculated with FAN-C. In order to define TADs, insulation scores were also calculated separately in the 25kb resolution KR matrices of each of the replicates (using FAN-C and as described in ^86^ with a window size of 500kb). Again, correlation between insulation scores of both replicates was high (r=0.94, **Extended Data Fig. 8b**). Definitive boundaries and TADs were then calculated in a merged 25kb matrix with a window size of 500kb and using a boundary score cutoff of 1 (**Extended Data Table 9**). CTCF motifs and their relative orientations were mined inside merged ATAC-seq peaks between the anterior and posterior pectoral fin samples using Clover ^87^, MA0139.1 Jaspar PWM, PWM score threshold of 8) and overlapped with previously calculated boundaries.

### HiChIP

HiChIP assays were performed as previously described [Mumbach et al. 2016], with some modifications. Briefly, 10 fins of stg. 30 skate embryos were fixed in a final concentration of 1% PFA for 10’ at room temperature. Fixation was quenched with 1M glycine up to a concentration of 0.125M. The tissue was then resuspended in 5 ml cell lysis buffer and homogenized with a Douncer on ice. After the lysis, nuclei were pelleted by centrifuging at 2500 rcf, and washed in 500 μl of lysis buffer. Chromatin digestion and ligation, ChIP, tagmentation and library preparation were performed as previously described ^88^.

### HiChIP analysis

Paired-end reads from HiChIP experiments were aligned to the skate genome using the TADbit pipeline ^89^ with default settings. Briefly, duplicate reads were removed, DpnII restriction fragments were assigned to resulting read pairs, valid interactions were kept by filtering out unligated and self-ligated events and multiresolution interaction matrices were generated. Dangling end read pairs were used to create 1D signal bedfiles that are equivalent to those of ChIP-seq experiments. Coverage profiles were then generated in bedgraph format using the *bedtools genomecov* tool, and bedgraph to bigwig conversions were also performed for visualization using the *bedGraphToBigWig* tool from *UCSC Kent Utils*. 1D signal bedgraph files were used to call peaks with MACS2 ^90^ using the no model and extsize 147 parameters and a FDR < 0.01.

FitHiChIP ^91^ was used to identify “peak-to-all” interactions at 10 kb resolution using HiChIP filtered pairs and peaks derived from dangling ends. Loops were called using a genomic distance between 20 kb and 2 Mb, and coverage bias correction was performed to achieve normalization. FitHiChIP loops with q-values smaller than 0.1 were kept for further analyses. Further filterings were performed to enrich enhancer-promoter interactions. First, loops established by two H3K4me3 peaks (likely promoter-promoter interactions) or no H3K4me3 peaks (likely enhancer-enhancer and others) were filtered out. Second, loops related to the H3K4me3 peak of the same gene promoter are grouped together into a common ‘regulatory landscape’, composed of a promoter anchor and several distal anchors. Then, regulatory landscapes with only one distal anchor were filtered out. Third, in order to filter out further spurious interactions we used the rationale that genomic bins that interact with a given promoter rarely do so in isolation. Therefore, we calculated a distance cutoff for ‘interaction gaps’ in regulatory landscapes. Regulatory landscapes containing interaction gaps bigger than the distance cutoff were trimmed and the distal anchors beyond the interaction gap discarded. The cutoff was determined for each sample independently by calculating the distribution of biggest gaps (calculating the biggest gap for each regulatory landscape) and setting the cutoff to the sum of the third quartile plus twice the interquartile range (classic outlier definition). Overlaps with ATAC-seq peaks in the pectoral fin were calculated using *bedtools intersect* (**Extended Data Fig. 9a**). Inter-TAD loops were also calculated using *bedtools intersect -c* using the TADs and the loops. Loops intersecting more than one TAD were considered inter-TAD loops. Randomized controls were generated shuffling TAD positions before the intersection using *bedtools shuffle*. For differential analysis between the anterior and the posterior fin, filtered distal anchors were fused when closer than 20kb using GenomicRanges *reduce* ^92^. The loops with the merged distal anchors are provided in the **Extended Data Table 10** and the loop numbers are displayed in the Venn Diagram in **Extended Data Fig. 9e**. In order to perform the differential analysis, the number of reads supporting the union set of loops was extracted for each of the sample replicates. Correlations shown in **Extended Data Fig. 9f, g** and the differential analysis performed using EdgeR ^93^ were calculated with this table (see **Supplementary Table 10**). An FDR cutoff of 0.1 was chosen to consider a loop to be significantly stronger in either the anterior or the posterior fin. Custom code used for enhancer-promoter loop filtering and differential analysis is included in the gitlab repository https://github.com/skategenome.

### Microsyntenic pair analysis

The analysis of microsyntenic pairs shared across the gnathostome lineage was based on the analysis described in ^94^. Briefly, we used the genome assembly and annotation presented in this work for the little skate in combination with public assemblies and annotations for mouse and garfish downloaded from ensembl (www.ensembl.org, *Mus musculus*: GRCm38v101, *Lepisosteus oculatus*: LepOcu1v104). Annotations in gtf format were converted to genepred with *gtfToGenePred* (https://github.com/ucscGenomeBrowser). Then, for each pair of consecutive genes in skates, we determined whether the ortholog pairs of genes in mouse and garfish were also consecutive (allowing 4 intervening gene models as in ^94^). Intergenic space between pairs of genes categorized as syntenic and non-syntenic in skates were overlapped with TAD boundaries and with TADs again using *bedtools intersect*. TADs were categorized according to the presence or absence of conserved microsyntenic pairs and then the overlap between the different TADs with ATAC-seq peaks or HiChIP loops was calculated again with *bedtools intersect*. List of conserved microsyntenic pairs is available in **Extended Data Table 11** and the code is available in the gitlab repository https://github.com/skategenome.

### TAD rearrangements in the skate lineage

In order to identify skate specific TAD rearrangements, global alignments were performed with *lastz* ^*95*^ against six different gnathostome genomes using as a reference the little skate assembly presented in this study. The chosen species were the thorny skate *Amblyraja radiata*, two species of shark (the white shark *Carcarodon carcarias* and the white-spotted bamboo shark *Chiloscyllium plagiosum*), one chimera (the elephant shark *Callorhinchus milii*) and a bony fish (the spotted gar *Lepisosteus oculatus*).The parameters of *lastz* were adapted to the phylogenetic distance with skate according to previous recommendations (^96^, see assemblies, substitution matrices and *lastz* parameters used in **Extended Data Table 12**). Then, syntenic chains and nets were devised as proposed elsewhere ^97^ and further polished with *chainCleaner* ^*98*^. Synteny breaks were then defined as the junctions between syntenic nets of any level, excluding those that were caused by the end of a scaffold for such genome assemblies that were not chromosome grade (white shark, elephant shark). The overlap between synteny breaks of different species was inferred using *bedtools multiinter*. Breaks that were found to be common in sharks, chimeras and a bony fish (garfish) were further considered. Then, distance between candidate synteny breaks and TAD boundaries (see **Extended Data Table 9**) was determined using *bedtools closest -d* and breaks that were located closer than 50 kb to a TAD boundary were discarded. Randomized analysis of overlap between synteny breaks and TAD boundaries (**Fig. 4b**) was carried out combining *bedtools closest* and *bedtools shuffle*. Finally, we selected candidate genes that displayed enhancer-promoter HiChIP interactions in the anterior or the posterior pectoral fin samples that crossed the synteny break, using again *bedtools intersect*. Enrichment of signalling pathways of candidate genes was performed using ReactomePA ^99^ and ClusterProfiler ^100^ R packages. Final synteny breaks and candidate genes are found in **Extended Data Table 13**, and the exact code used is found in the code repository https://github.com/skategenome.

### Whole mount *in situ* hybridization

Skate and shark embryos were recovered from egg cases at stage 27 and 30 and fixed by 4% PFA at 4ºC overnight. Next day, the embryos were rinsed by PBS-0.1% Tween 3 times, soaked in 100% MeOH, and stored -80ºC. Whole mount in situ hybridization was conducted as previously described except for hybridizing the embryos and probes at 72ºC ^3^.

### Real time PCR

The pectoral fins of three shark juveniles (*Scyliorhinus retifer*) were dissected out in DEPC-PBS at stage 30. Three replicates were prepared. Total RNA was separately extracted from each replicate by Trizol (Invitrogen). cDNA was synthesized from the total RNA by iScript™ cDNA Synthesis Kit (Bio-Rad). Then, real-time PCR for *gapdh* and *prickle1* was conducted with KAPA HiFi HotStart ReadyMix PCR Kit (Kapa Biosystems) and Applied Biosystems 7300 Real time PCR system. The primers used in this study are listed in **Extended Data Table 7**. The obtained Ct value from RT-PCR was converted to arbitrary gene expression values.

### Cell elongation analysis

Pectoral fins were dissected from stage 29 skate embryos and fixed by 4% paraformaldehyde overnight. Next day, the fins were rinsed by PBS including 0.1% Triton X-100 and incubated in the blocking buffer (10 % sheep serum and 0.1% BSA in PBS-0.1% Triton X-100) at room temperature for an hour. Then, the blocking buffer was replaced by the blocking buffer, including CellMask Deep Red Plasma membrane Stain (1/1000 dilution, Invitrogen) and DAPI (1:4000 dilution), and incubated at 4ºC overnight. Subsequently, the fins were washed for 1 hour five times by PBS-Triton-X100 and mounted on glass slides. The fins were then scanned by a confocal microscope (ZEISS, LSM510 META). The scanned images were incorporated into Fiji and cell outlines in fin mesenchyme were manually traced. The cell elongation ratio was automatically calculated by the macro “Tissue Cell Geometry Stats” included in Fiji.

### ROCK inhibitor treatment

To test the function of the PCP pathway in the pectoral fin development, skate embryos were treated by Y27632, a ROCK inhibitor, from stage 29 to stage 31 and investigated for their fin morphology. 500 μl of ROCK inhibitor (stock 50 mM, final 50 μM, Selleck chemicals) or DMSO solution (negative control) was added to 500 ml of artificial saltwater (Instant Ocean), and five skate embryos at stage 29 for each condition were kept submerged in these solutions. Once the negative control embryos reached stage 31, all embryos were fixed by 4% PFA and their total body length was measured under a stereomicroscope. *N*=5 / condition.

### Morphometrics analysis of skate fins

Skate embryos at each stage were photographed from the ventral side. A landmark scheme was designed to capture the shape of the pectoral fin (**Extended Data Fig. 13e**). Six homologous landmarks and three curves were assessed in each specimen; curves were used to generate sliding semi-landmarks. Specimens were digitized in R using the package Stereomorph ^101^. Digitized files were then uploaded to ShinyGM ^102,103^, where all downstream analyses were performed. Specimens were aligned using a Generalized Procrustes Analysis to account for shape differences due to differences in specimen size, specimen orientation, and scaling. A morphospace was generated using these aligned landmark coordinates; deformation grids were generated for the control stage 31 and ROCK-inhibited stage 31 specimens (**Fig. 4f**). A linear model was run to test for the impact of length, treatment, and stage on shape. Both treatment and stage were significantly associated with shape (p = 0.002 and p = 0.001, respectively); as expected, total length was not significantly associated with the size-corrected shapes (p = 0.711).

### Transgenic enhancer activity assay

Shark and skate HoxA2-A1 enhancers were cloned into pCR™8/GW/TOPO® vector (Invitrogen) by PCR. The primers are listed in **Extended Data Table 7**. The cloned enhancers were transferred into pXIG-cfos-EGFP vector by Gateway™ LR Clonase™ II (Invitrogen)^55^. The created vectors were injected into one-cell stage zebrafish eggs with *Tol2* mRNA as previously described ^104^. The injected embryos were observed under a stereo-type fluorescent microscope and photographed at 48 hours post fertilization.

### Phylome reconstruction

The phylome of *Leucoraja erinacea*, meaning the collection of phylogenetic trees for each protein-coding gene in its genome, was reconstructed using an automated pipeline that mimics the steps one would take to build a phylogenetic tree and based on the PhylomeDB pipeline ^105^. First a database with the proteomes (i.e. full set of protein-coding genes) of 21 species was built that included *Leucoraja erinacea* (see table S1 for a full list of species included). Then a blastp search was performed against this database starting from each of the proteins included in the *L. erinacea* genome. Blast results were filtered using an e-value threshold of 1e-05 and a query sequence overlap threshold of 50%. The number of hits was limited to the best 250 hits for each protein. Then a multiple sequence alignment was performed for each set of homologous sequences. Three different programs were used to build the alignments (Muscle v3.8.1551 ^106^, mafft v7.407 ^107^ and kalign v2.04 ^108^) and the alignments were performed in forward and in reverse resulting in six different alignments. From this group of alignments a consensus alignment was obtained using M-coffee from the T-coffee package v12.0 ^109^. Alignments were then trimmed using trimAl v1.4.rev15 (consistency-score cut-off 0.1667, gap-score cut-off 0.9) ^110^. IQTREE v1.6.9 ^111^ was then used to reconstruct a maximum likelihood phylogenetic tree. Model selection was limited to 5 models (DCmut, JTTDCMut, LG, WAG, VT) with freerate categories set to vary between 4 and 10. The best model according to the BIC criterion was used. 1000 rapid bootstrap replicates were calculated. A second phylome starting from *Danio rerio* was also reconstructed following the same approach. All trees and alignments are stored in phylomedb ^105^ with phylomeIDs 247 for the *L. erinacea* phylome and 275 for the *D. rerio* phylome (http://phylomedb.org).

### Species tree reconstruction

A species tree was reconstructed using a gene concatenation approach. The trimmed alignments of 102 protein families with a single ortholog per species were concatenated into a single multiple sequence alignment. IQTREE ^111^ was then used to reconstruct the species tree using the same parameters as above. The final alignment contained 48,958 positions. The model selected for tree reconstruction was JTTDCMut+F+R5. Additionally, duptree ^112^ was used to reconstruct a second species tree using a super tree method. Duptree searches for the species tree that minimizes the number of duplications inferred when each gene is reconciled with the species tree. All trees built during the phylome reconstruction process were used to reconstruct this species tree. The two topologies were fully congruent.

### Skate MethylC-seq library preparation

MethylC-seq library preparation was performed as described previously ^113^. Briefly, 1000 ng of genomic DNA extracted from the embryonic stage31 and adult skate pelvic and pectoral fins was spiked with unmethylated λ phage DNA (Promega). DNA was sonicated to ∼300 bp fragments using M220 focused ultrasonicator (Covaris) with the following parameters: peak incident power, 50W; duty factor, 20%; cycles per burst, 200; treatment time, 75 sec. Sonicated DNA was then purified, end-repaired using End-It™ DNA End-Repair Kit (Lucigen) and A-tailed using Klenow Fragment (3’→5’ exo-) (New England Biolabs) followed by the ligation of NEXTFLEX^®^ Bisulfite-Seq Adapters. Bisulfite conversion of adaptor-ligated DNA was performed using EZ DNA Methylation-Gold Kit (Zymo Research). Library amplification was performed using KAPA HiFi HotStart Uracil+ DNA polymerase (Kapa Biosystems). Library size was determined by the Agilent 4200 Tapestation system. The libraries were quantified using the KAPA library quantification kit (Roche).

### Skate methylome data analysis

Embryonic stage31 and adult skate pelvic and pectoral fin DNA methylome libraries were sequenced on the Illumina HiSeq X platform (150 bp, PE). Elephant shark *Callorhinchus milii* raw whole genome bisulphite sequencing data (adult liver) was downloaded from PRJNA379367 ^114^. Zebrafish *Danio rerio* raw whole genome bisulphite sequencing data (adult liver) was downloaded from GSE122723 ^115^. Sequenced reads in FASTQ format were trimmed using the Trimmomatic software (ILLUMINACLIP:adapter.fa:2:30:10 SLIDINGWINDOW:5:20 LEADING:3 TRAILING:3 MINLEN:50). Trimmed reads were mapped to the Leri_hhj.fasta genome reference (containing the lambda genome as chrLambda) using WALT ^116^ with the following settings: -m 10 -t 24 -N 10000000 -L 2000. Mapped reads in SAM format were converted to BAM format; BAM files were sorted and indexed using SAMtools ^117^. Duplicate reads were removed using Picard Tools v2.3.0 (http://broadinstitute.github.io/picard/). Genotype and methylation bias correction were performed using MethylDackel (MethylDackel extract Leri_hhj_lambda.fasta $input_bam -o $output --mergeContext --minOppositeDepth 5 --maxVariantFrac 0.5 --OT 10,110,10,110 --OB 40,140,40,140) (https://github.com/dpryan79/MethylDackel). Methylated and unmethylated calls at each genomic CpG position were determined using MethylDackel (MethylDackel extract Leri_hhj_lambda.fasta $input_bam -o output –mergeContext). DNA methylation profiles at differentially accessible ATAC-seq peaks between embryonic pelvic and pectoral fin samples was performed using deepTools2 *computeMatrix reference-*point and *plotHeatmap* ^*118*^.

## ACKNOWLEDGEMENTS

We thank Dr Richard Schneider, Prof. David Sherwood, and the Marine Biological Laboratory Embryology course for provision of lab space, Louise Bertrand and Leica Microsystems for microscopy support, and David Remsen, Scott Bennett, Dan Calzarette, and the staff of the Marine Biological Laboratory and MBL Marine Resources Center for technical and animal husbandry assistance. We thank Andrew Gillis for support and advice with RNA-seq and skate functional experiments.

T.N., D.N., and A.A. were supported by institutional support provided by the Rutgers University School of Arts and Sciences and the Human Genetics Institute of New Jersey and a Marine Biological Laboratory research grant. DN was further supported by the NIH-IRACDA funded INSPIRE program at Rutgers University. F.M. and D.S.R. were supported by funding from the Okinawa Institute for Science and Technology. D.S.R is grateful for the support of the Marthella Foskett-Brown Chair in Computational Biology. D.G.L. and R.D.A. were supported by a grant from the Deutsche Forschungsgemeinschaft (LU 242672-1) and by a Helmholtz ERC Recognition Award grant from the Helmholtz-Gemeinschaft (ERC-RA1045 0033). R.D.A. and C.P. were supported by EMBO Postdoctoral Fellowships (EMBO ALTF 537-2020 and ALTF 346-2020, respectively). J.J.T. and J.L.G.S. were supported by the European Research Council (ERC, grant agreement No 740041) and the Spanish Ministerio de Economía y Competitividad (grant PID2019-103921GB-I00). F.M. is supported by the Royal Society (URF\R1\191161). V.A.S. was supported by a Wolfson College Junior Research Fellowship and Marine Biological Laboratory Whitman Early Career Fellowship. J.L.-R. is supported by the Spanish Ministerio de Ciencia e Innovacion (PID2020-113497GB-I00). A.V. and F.D. were supported by NIH grants R01DE028599 and R01HG003988. Research conducted at the E.O. Lawrence Berkeley National Laboratory was performed under US Department of Energy contract DE-AC02-05CH11231, University of California.

## AUTHOR CONTRIBUTIONS

J.L.G.S., F.M., J.J.T., D.S.R, and D.G.L. conceived the study and designed the experiments. F.M., E.D.L.C.M., N.S. and J.L.G.S coordinated the sequencing of the little skate genome and F.M. assembled and annotated the genome. R.D.A., P.M.M.G., L.Y., J.H.G, J.D. and D.G.L performed analyses on 3D chromatin organization. F.M., D.S.R designed and performed synteny and comparative analyses. M.M.H., F.M. and T.G. performed phylogenetic and phylogenomic analyses. E.D.L.C.M., C.P., S.N., R.D.A., J.J.T., I.C., L.G.F., I.S. and J.L.R. performed and analysed transgenics, ATAC-seq and RNA-seq experiments. V.A.S and C.H. performed and analysed additional RNA-seq experiments. F.D. and A.V. performed additional functional assays. K.S., P.E.D., A.G.R. and O.B. performed and analyzed DNA methylation experiments. D.N., A.A., and T.N. conducted embryonic experiments of skate and sharks. J.L.G.S., J.J.T., F.M., D.S.R., and D.G.L. wrote the manuscript with input from all authors.

## ETHICS DECLARATIONS

Competing interests

The authors declare no competing interests.

## Notes

### Competing Interest Statement

The authors have declared no competing interest.

