## Extended Data for "The little skate genome and the evolutionary emergence of wing-like fin appendages"

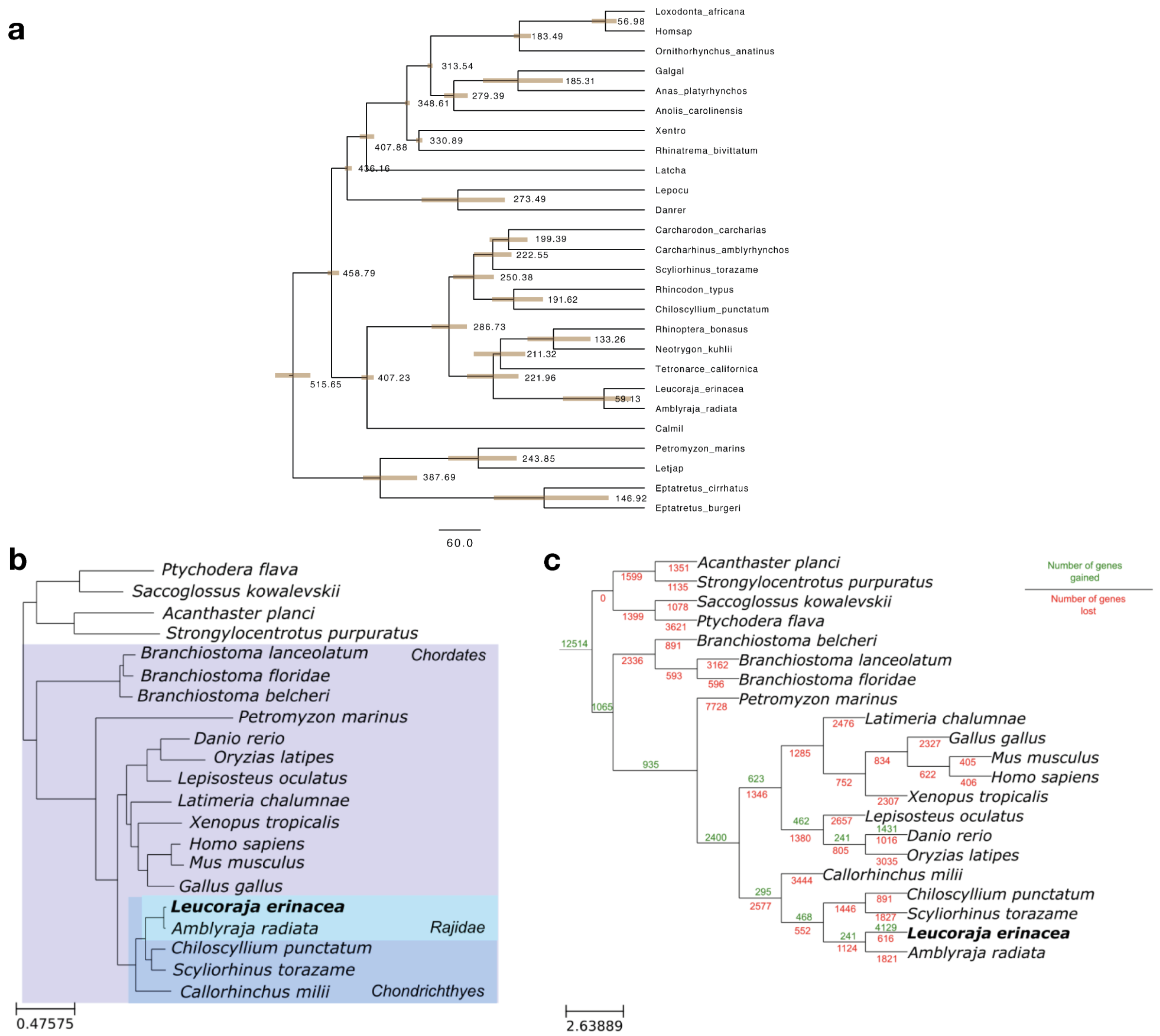


**Extended Data Figure 1. Extended chronogram and gene gains and loss from Phylome. a**, Chronogram of selected vertebrate species based on 74145 aminoacid position analyzed using Phylobayes assuming the CIR model with birth-death priors and and soft-bounds on divergence times (calibrations from [1]). **b**, Species using for Phylome reconstruction and **c,** Gain and loss of genes along the branch as inferred using the Phylome pipeline.

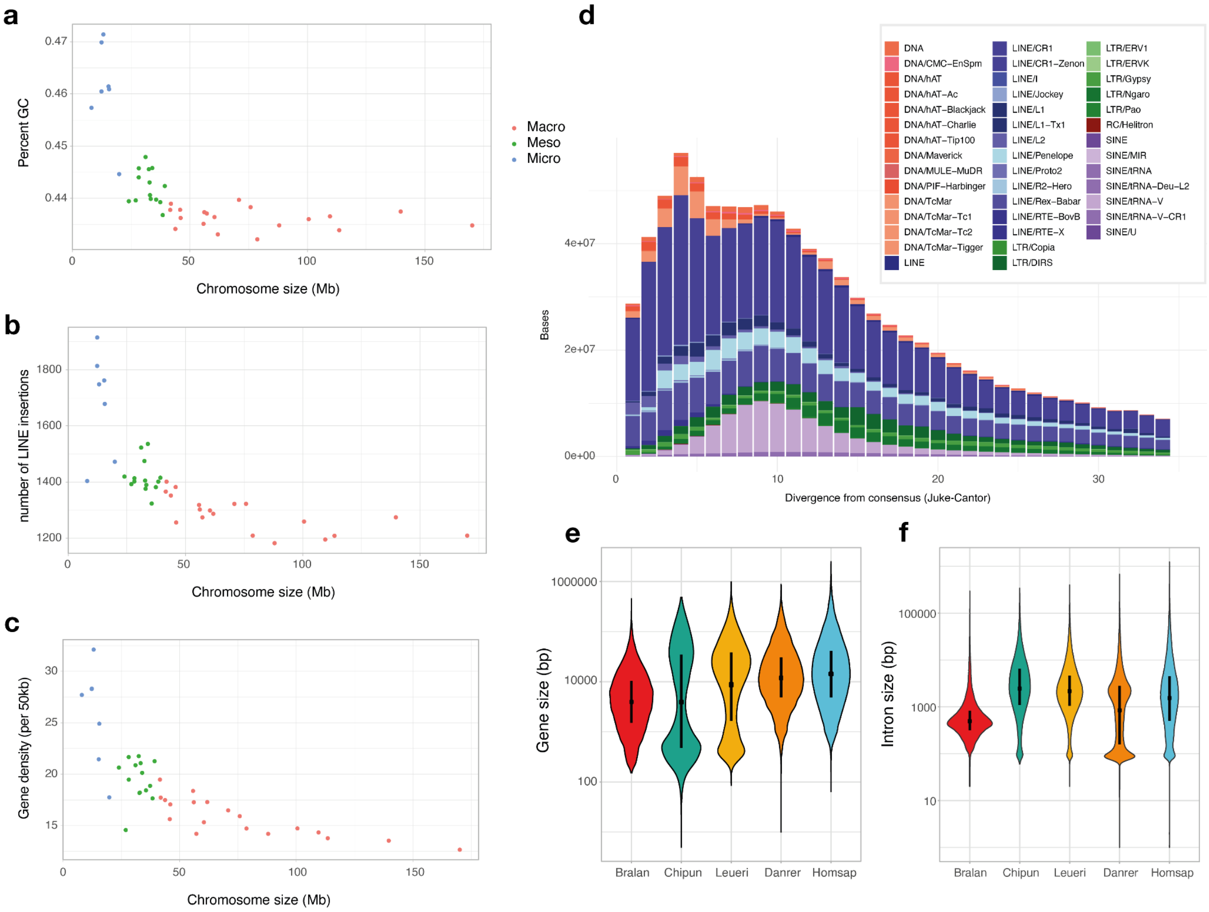


**Extended Data Fig. 2. Characteristics of skate chromosomes, repeat content and skate genome. a-c,** Characteristics and classification of skate chromosomes according to their size (x-axis) and GC% (a), number of LINE insertions (b) and gene density (c) per 50kb window. **d,** Repetitive landscape computed as JC divergence of repeat occurrence toward the consensus element in the repeat library. **e-f**, Distribution of gene and intron size in selected chordate species: amphioxus (*Branchiostoma floridae*, Bralan), the cloudy catshark (*Chiloscyllium punctatum*, Chipun), the little skate (*Leuraja erinacea*, Leueri), the zebrafish (*Danio rerio*, Danrer) and human (*Homo sapiens*, Homsap).

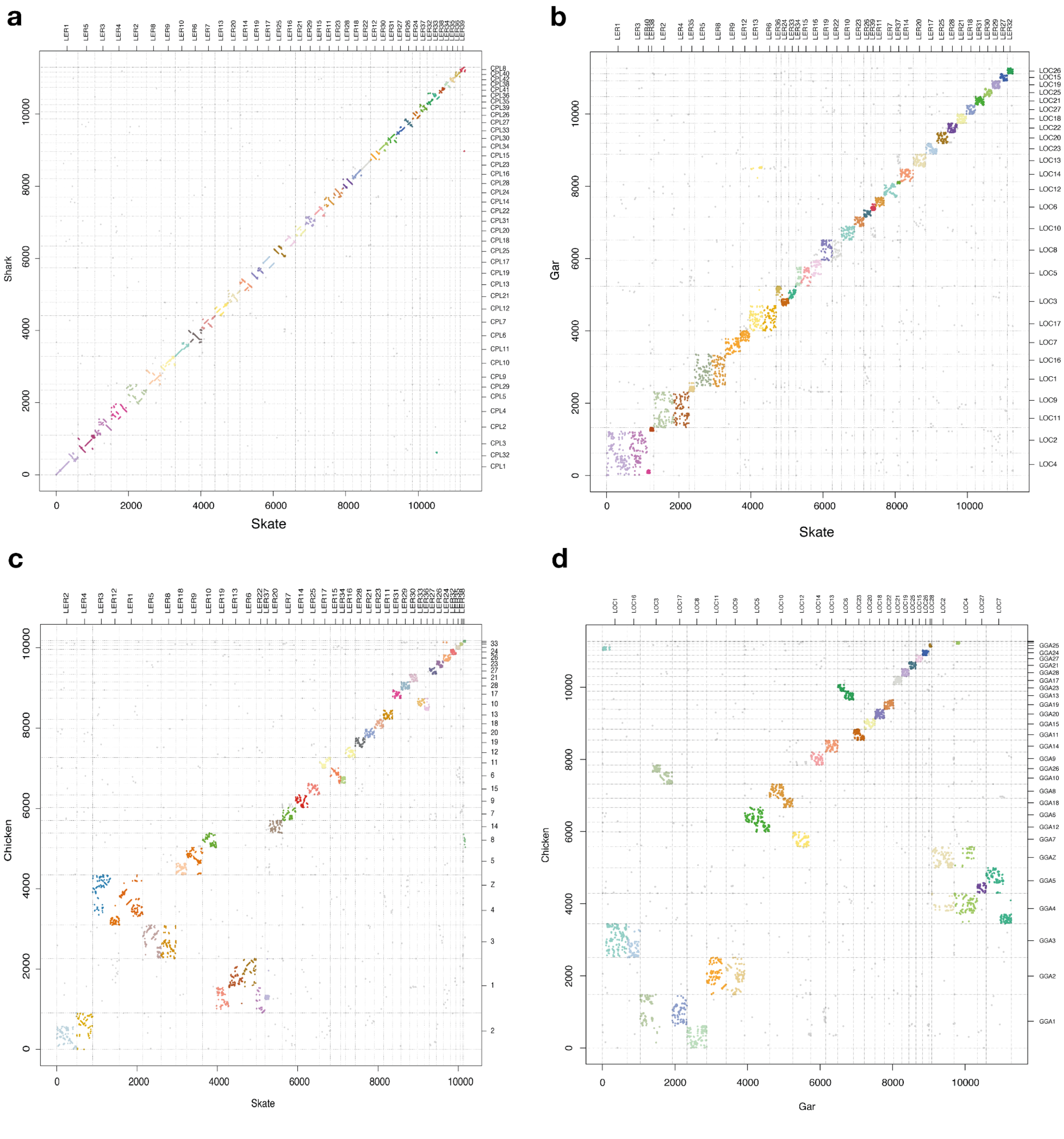


**Extended Data Fig 3. Synteny and disruption of gene order. a-c,** Synteny plots between the little skate (*Leucoraja erinacea*) and selected vertebrate species: **a,** white–spotted bamboo shark (*Chiloscyllium plagiosum*); **b**, gar (*Lepisosteus oculatus*); **c**, chicken (*Gallus gallus*). **d**, Synteny plot between gar and chicken.

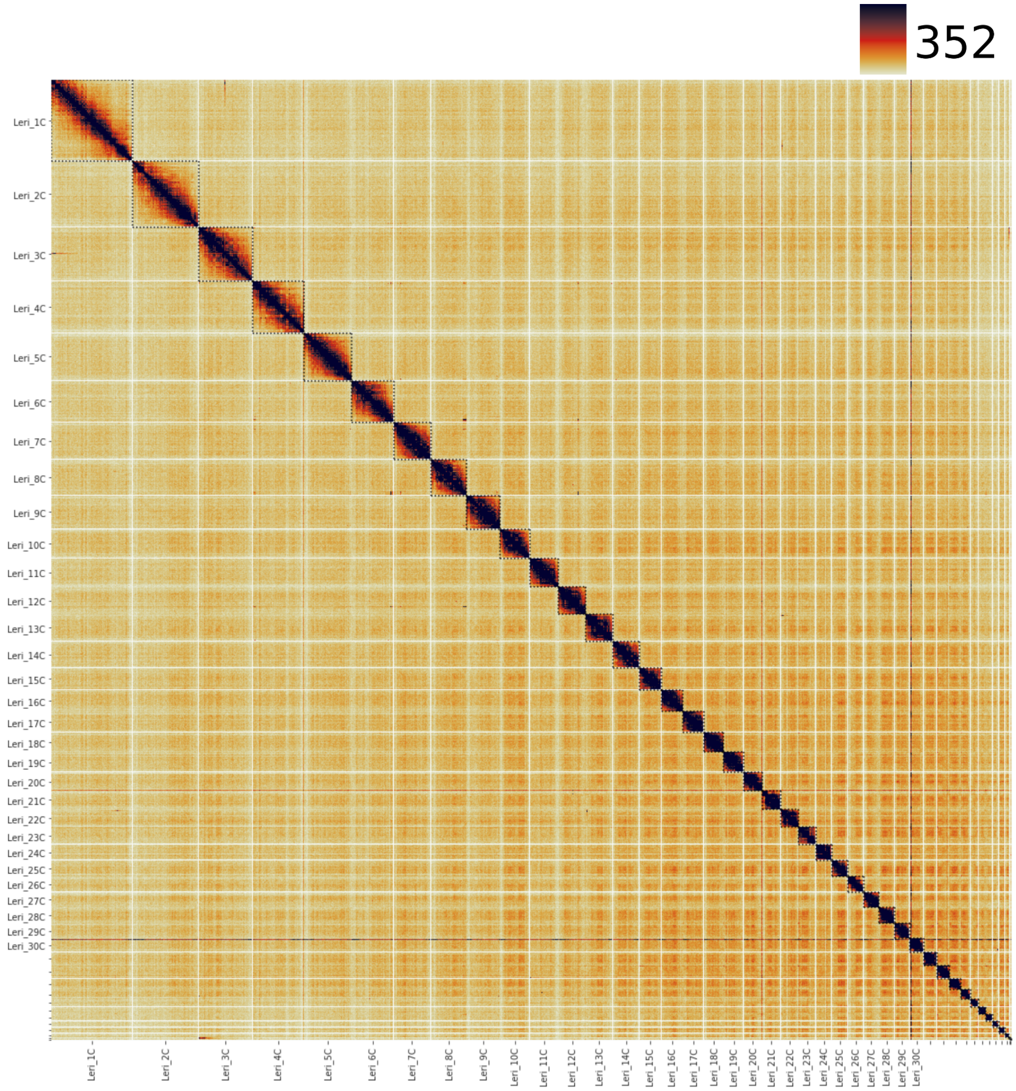


**Extended Data Figure 4: Skate chromosome territories.** Skates display a full set of condensins and therefore their genome is organized in chromosome territories. Hi-C map at 1Mb resolution of the entire *L. erinacea* genome. Chromosome ends are delineated with dashed lines. Intrachromosomal contacts are enriched.

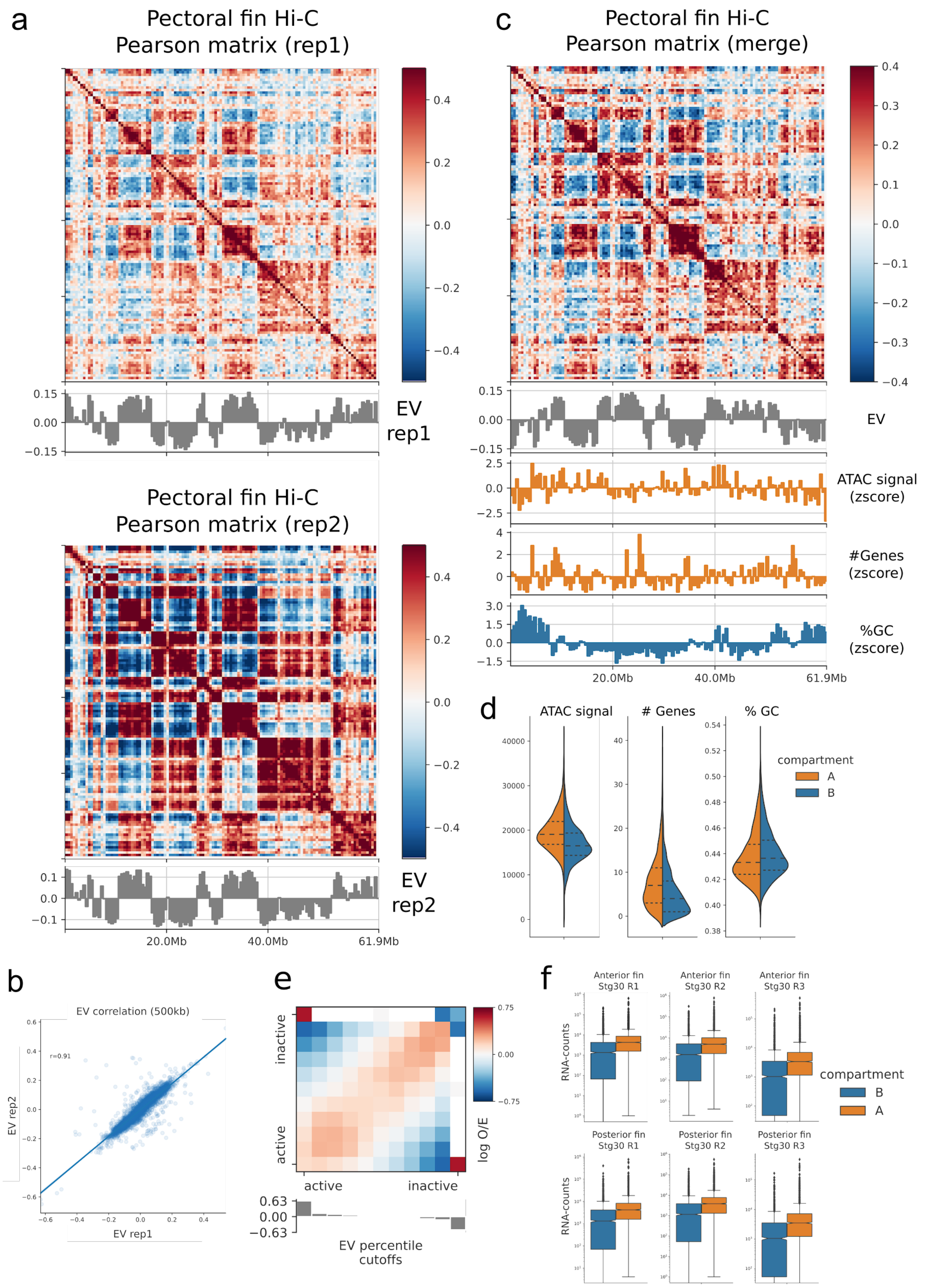

a
Pectoral fin Hi-C
Pearson matrix (rep1)
EV
rep1
Pectoral fin Hi-C
Pearson matrix (rep2)
EV
rep2
20.0Mb
40.0Mb
61.9Mb
c
Pectoral fin Hi-C
Pearson matrix (merge)
EV
ATAC signal
(zscore)
#Genes
(zscore)
%GC
(zscore)
d
ATAC signal
### Genes
% GC
compartment
A
B
b
EV correlation (500kb)
r=0.91
EV rep2
EV rep1
e
inactive
active
log O/E
EV percentile
cutoffs
f
Anterior fin
Stg30 R1
Anterior fin
Stg30 R2
Anterior fin
Stg30 R3
Posterior fin
Stg30 R1
Posterior fin
Stg30 R2
Posterior fin
Stg30 R3
RNA-counts
compartment
B
A

**Extended Data Figure 5. The skate genome is organized in A/B compartments. a.** 500 kb resolution Pearson matrices of a representative chromosome (Leri_11C) and their associated eigenvectors showing marked compartmentalization in A/B compartments in both replicates. **b.** Eigenvector correlation among the two replicates. **c.** Merged Pearson matrix presented together with its eigenvector, the normalized signal for ATAC-seq in anterior pectoral fin, the number of gene models and the percentage of GC content. As shown in **d**, the A compartment in skates correlates with chromatin accessibility and the number of gene models, but no clear correlation was observed with the GC content. **e.** Saddle plot demonstrating the aggregated enrichment of homotypic A-A and B-B interactions. **f.** Gene expression in either the A or the B compartment as measured with bulk RNA-seq performed in the anterior and posterior portions of the skate pectoral fin at the equivalent stage (Stg30, top: anterior, bottom: posterior).

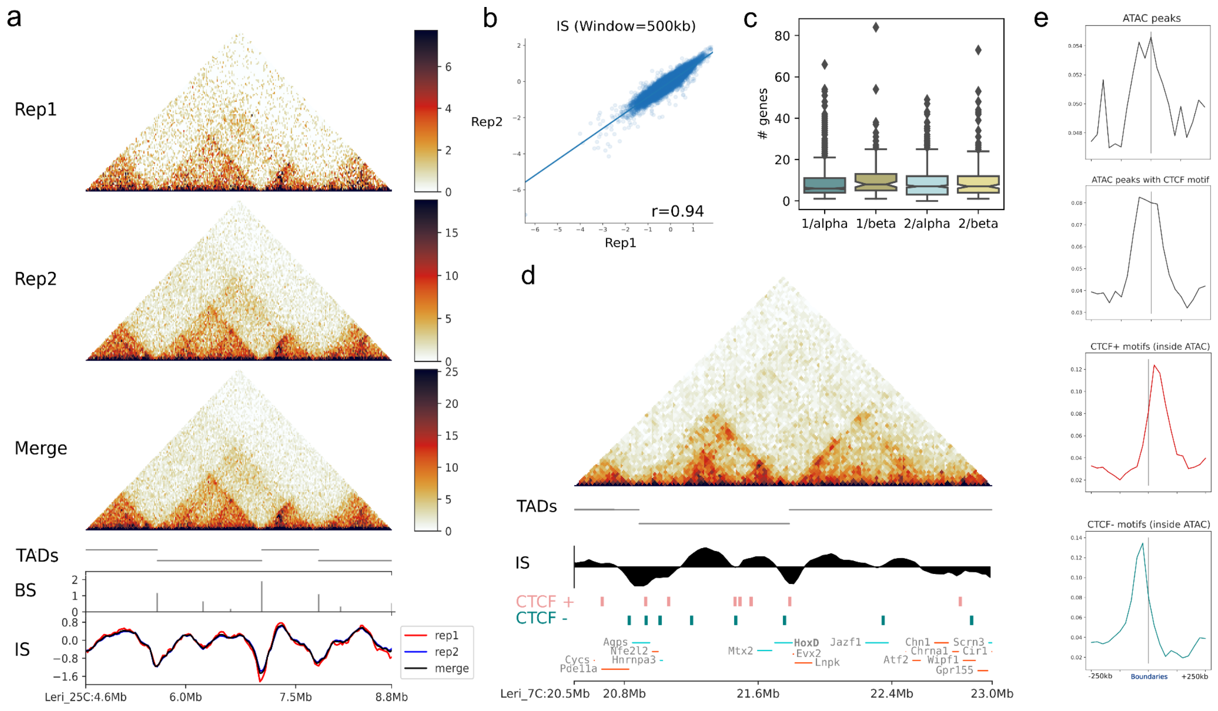


**Extended Data Figure 6. Skate chromosomes are organized in TADs flanked by convergent CTCF sites. a.** Hi-C interaction matrices in skate pectoral fins in either of the two replicates and the merge (25kb resolution). The TAD calling performed in the merged matrix and the associated boundary scores (BS) and insulation scores (IS) are shown below (window size of 500kb). **b.** Insulation score correlations between the two replicates. **c.** Gene content of TADs associated to the different paralogous segments of the genome originated after the two rounds of WGD (1 or 2 for the 1R, alpha or beta for the 2R). **d.** Hi-C matrix around the HoxD locus showing the conserved bipartite configuration in two TADs with HoxD genes located precisely at the boundary. TADs, insulation scores and ATAC-seq peaks containing the CTCF motif are shown. The tendency of having divergent CTCF sites at insulation minima is observable. **e.** From top to bottom, enrichment around TAD boundaries (+-250kb) of ATAC-seq peaks and ATAC-seq peaks containing the CTCF motif regardless of the strand, in the plus and in the minus strand.

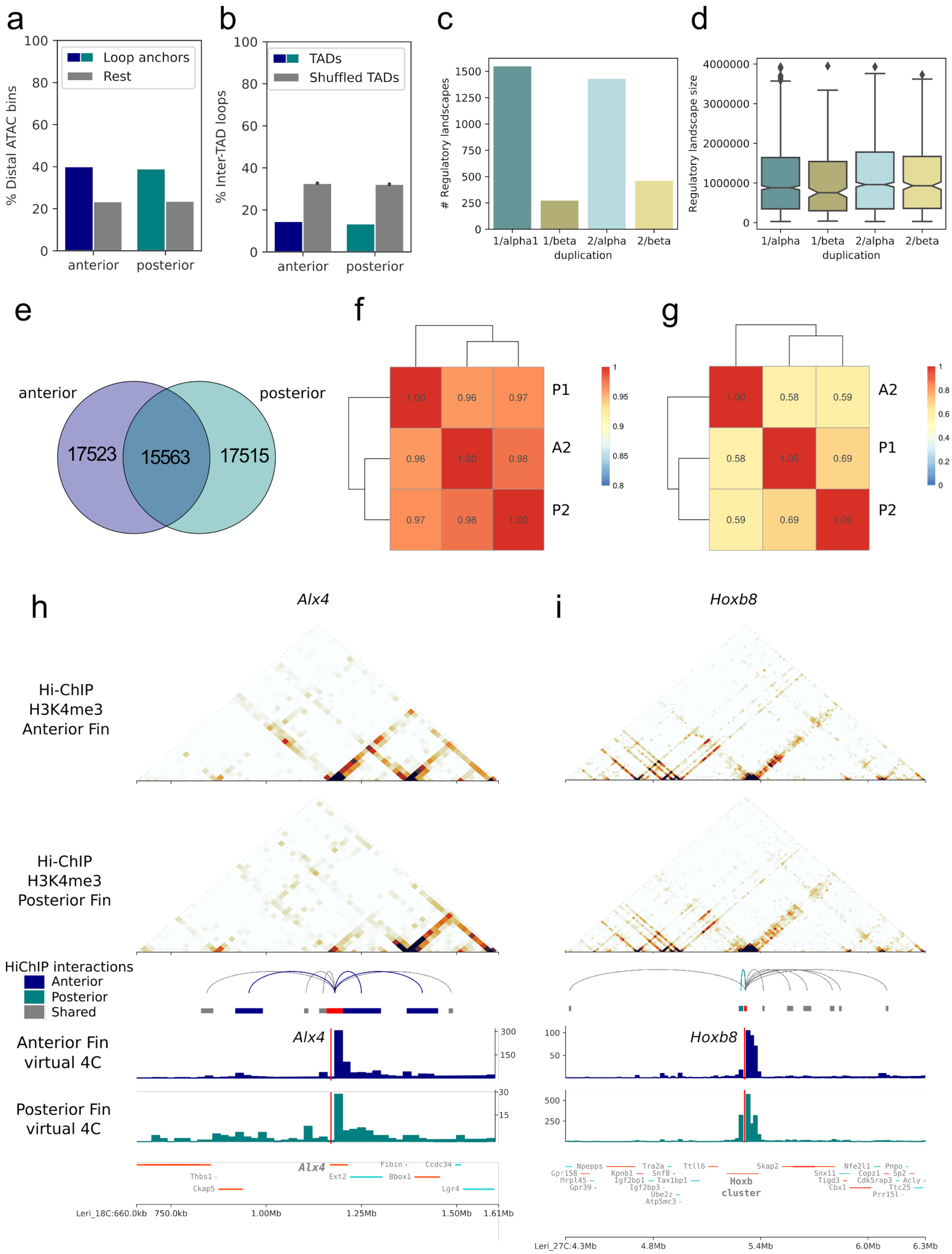

a
% Distal ATAC bins
Loop anchors
Rest
anterior
posterior
b
% Inter-TAD loops
TADs
Shuffled TADs
anterior
posterior
c
### Regulatory landscapes
1/alpha1
1/beta
2/alpha
2/beta
duplication
d
Regulatory landscape size
1/alpha
1/beta
2/alpha
2/beta
duplication
e
anterior
posterior
17523
15563
17515
f
P1
A2
P2
g
A2
P1
P2
h
Alx4
Hi-ChIP H3K4me3 Anterior Fin
Hi-ChIP H3K4me3 Posterior Fin
HiChIP interactions
Anterior
Posterior
Shared
Anterior Fin virtual 4C
Posterior Fin virtual 4C
Leri_18C:660.0kb
750.0kb
1.00Mb
1.25Mb
1.50Mb
1.61Mb
i
Hoxb8
Hoxb cluster
Leri_27C:4.3Mb
4.8Mb
5.4Mb
6.0Mb
6.3Mb

**Extended Data Figure 7. H3K4me3 HiChIP unveils the regulatory landscapes of active genes in the anterior and posterior portions of the skate pectoral fin. a.** Proportion of distal loop anchors that also correspond to distal ATAC-seq peaks in the pectoral fin in both the anterior and posterior H3K4me3 HiChIP datasets. **b.** Proportion of inter-TAD interactions calculated in the anterior and posterior HiChIP datasets compared to a random shuffling of the TADs (gray). **c.** Number of regulatory landscapes (defined as the group of interactions anchored by a single gene promoter) belonging to the different paralogous segments of the genome originated after the two rounds of WGD (1 or 2 for the 1R, alpha or beta for the 2R). **d.** Regulatory landscape sizes observed in the paralogous segments of **c** defined as the genomic space spanning from the two more distal loop anchors anchored to a given promoter. **e.** Venn diagram showing the overlap among loops identified in the anterior and posterior H3K4me3 HiChIP datasets. **f.** Pearson correlation of the three valid replicates (1 for anterior and 2 for posterior fins). The correlation between the matrices is limited to the non-redundant set of interactions (50,601, union set of the Venn diagram in **e**). **g**. Same as in **f** but using Spearman. **h.** Anterior specific contacts in the *Alx4* regulatory landscape (dark blue). **i.** Posterior specific contacts in the *Hoxb8* regulatory landscape (turquoise).

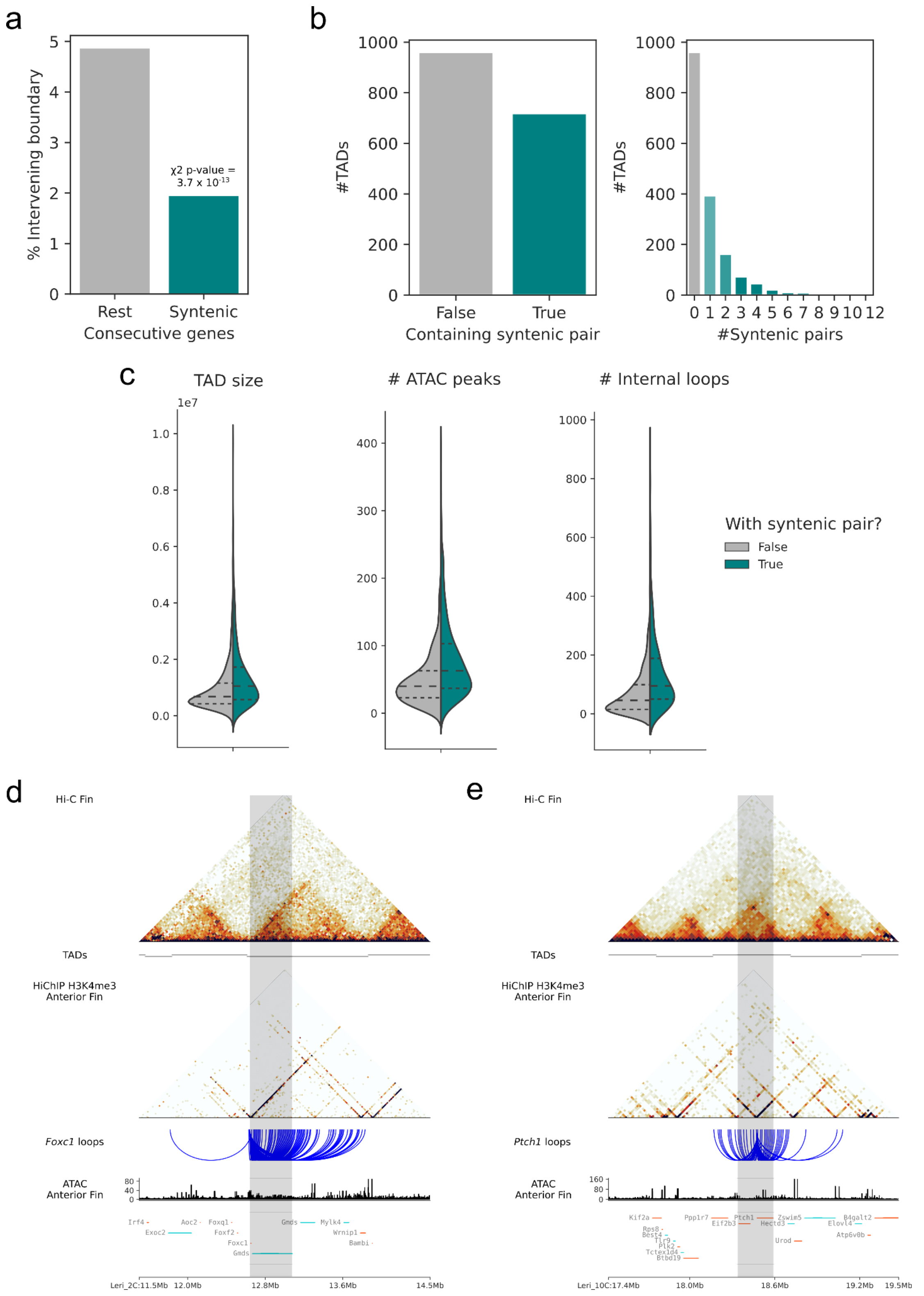

a
% Intervening boundary
χ2 p-value = 3.7 x 10-13
Rest
Syntenic
Consecutive genes
b
#TADs
False
True
Containing syntenic pair
#Syntenic pairs
c
TAD size
### ATAC peaks
### Internal loops
1e7
With syntenic pair?
False
True
d
Hi-C Fin
TADs
HiChIP H3K4me3 Anterior Fin
Foxc1 loops
ATAC Anterior Fin
Leri_2C:11.5Mb
12.0Mb
12.8Mb
13.6Mb
14.5Mb
e
Hi-C Fin
TADs
HiChIP H3K4me3 Anterior Fin
Ptch1 loops
ATAC Anterior Fin
Leri_10C:17.4Mb
18.0Mb
18.6Mb
19.2Mb
19.5Mb

**Extended Data Figure 8. Deeply conserved microsyntenic TADs in jawed vertebrates. a.** Intergenic spaces between microsyntenic pairs conserved across vertebrates (present in skate and osteichthyes, here mouse and garfish) are devoid of TAD boundaries. **b.** 40% of skate TADs contain a deeply conserved microsyntenic pair. Several of them contain more than one association. **c.** TADs containing deeply microsyntenic associations are bigger, contain more ATAC-seq peaks and more loops as defined using HiChIP. *Foxc1/Gmds* (**d**) and *Ptch1/Eif2b3* (**e**) are examples of deeply conserved microsyntenic associations. Microsyntenic area is shaded in grey. Hi-C, TADs, HiChIP and ATAC-seq data are shown along with the gene tracks.

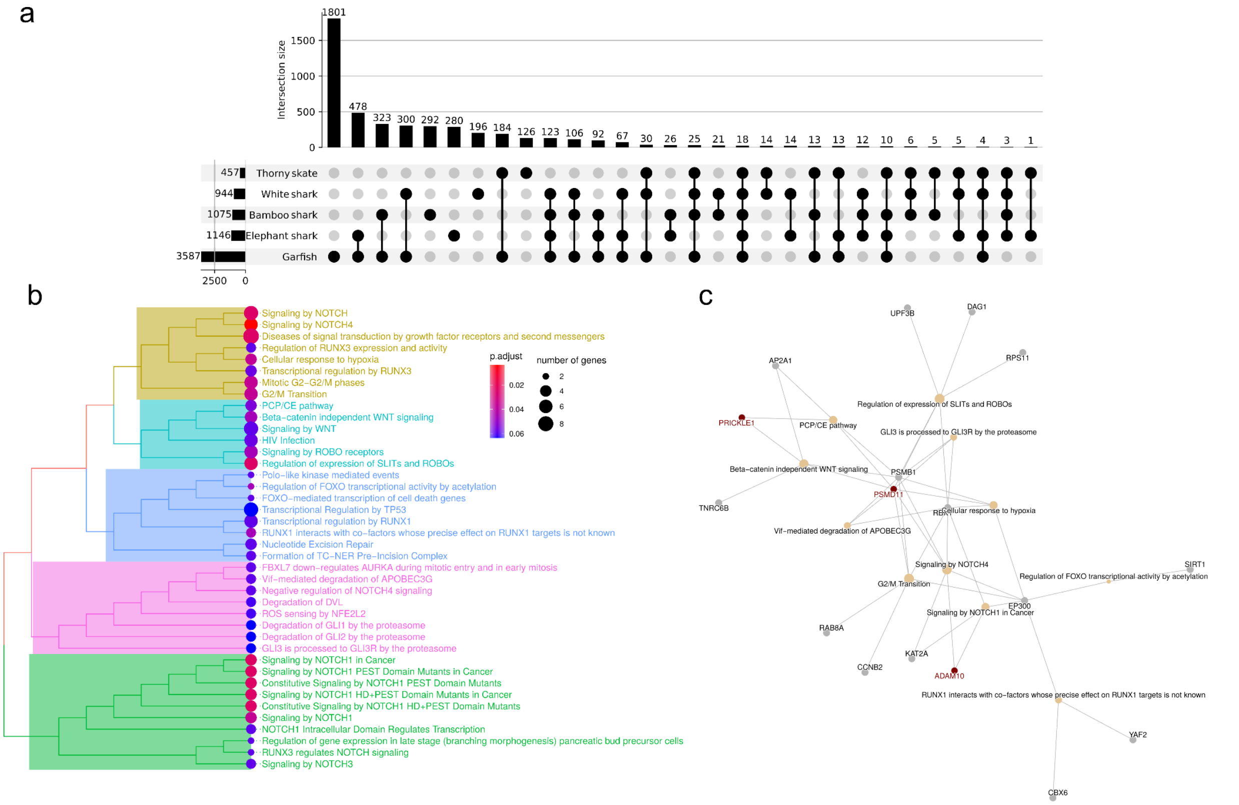


**Extended Data Figure 9. Rearranged TADs in the skate lineage involve PCP-related genes. a.** Extended version of the upset plot presented in **Fig. 4a** with the quantification of the synteny breaks found in the whole set of species combinations. **b.** ReactomePA [2] clustering of significant terms found in the set of candidate genes for regulatory rearrangements in the anterior pectoral fin. A selection of these terms is shown in **Fig. 4c**. **c.** Cnetplot showing the relationship of candidate genes with each of the different enriched terms.

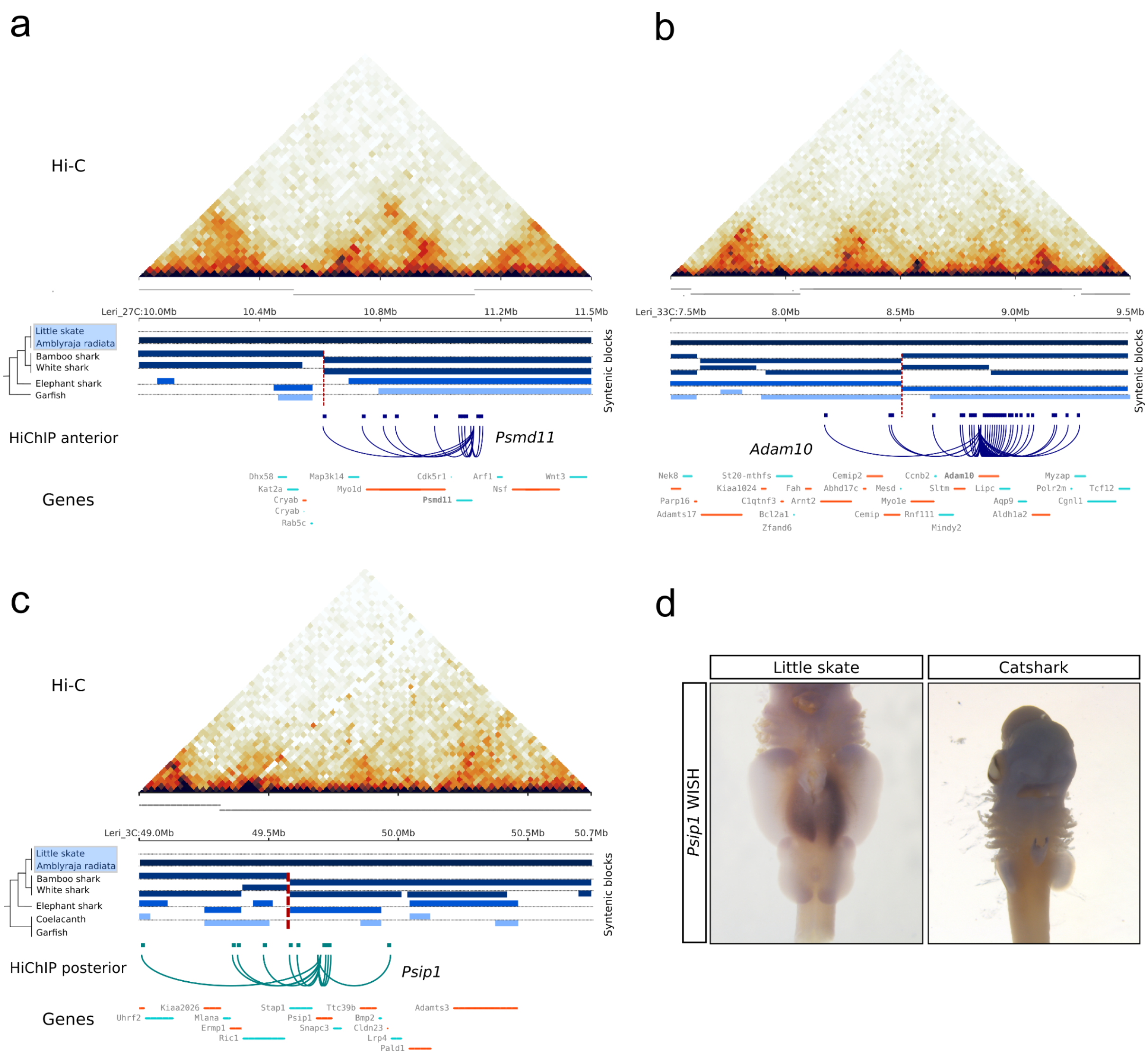


**Extended Data Figure 10. Examples of TAD rearrangements in the skate lineage. a.** Candidate rearrangement at the *Psmd11* locus, implicated in the PCP pathway. Pectoral fin Hi-C map is shown on top together with the TAD predictions. Below, the synteny blocks that are shared with the different species studied and the candidate synteny break is highlighted in red. Finally, arachnogram with the contacts devised from the anterior fin H3K4me3 HiChIP experiment. **b.** Same as in **a,** but for the Notch-signalling related gene *Adam10*. **c.** Same as in **a** and **b** but for the Hox activator *Psip1*. Note that this time the presented H3K4me3 HiChIP is from posterior pectoral fins. **d.** Whole mount *in situ* hybridization against *Psip1* in both the little skate *L.erinacea* and the catshark *S.retifer* shows species-specific expression of *Psip1* in the anterior portion of the skate pectoral fins.

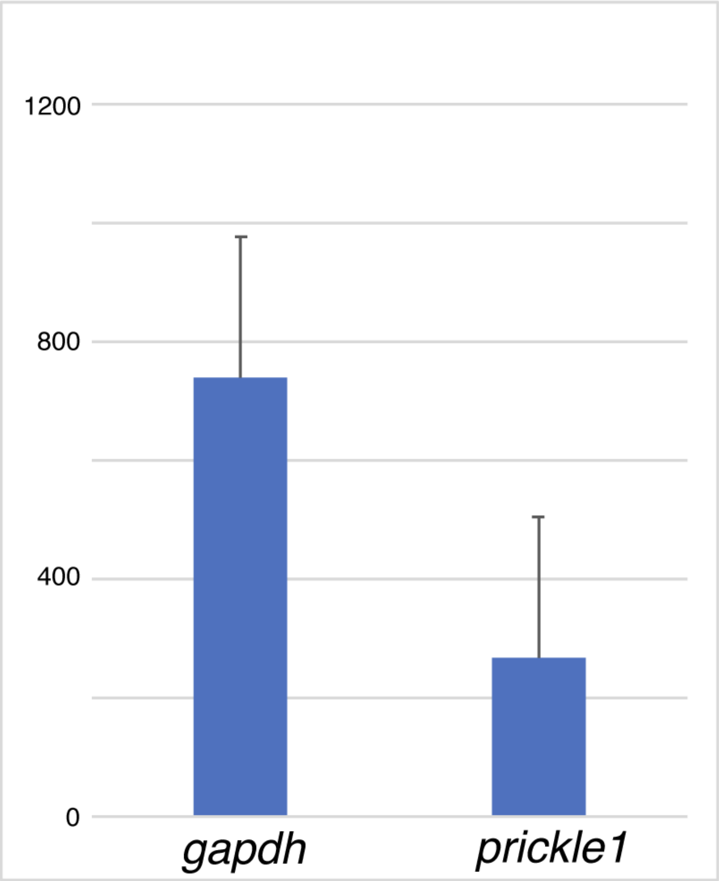


**Extended Data Figure 11. Real-time PCR of *gapdh* and *prickle 1* in shark pectoral fins.**
The transcripts of *gapdh* and *prickle1* in the shark pectoral fin (*Scyliorhinus retifer*) at stage 30 were detected by real-time PCR. Three replicates were performed. The vertical axis shows the arbitrary expression values calculated from Ct-values of the real-time PCR. The error bars are standard errors. Both *gapdh* and *prickle 1* transcripts were successfully detected.

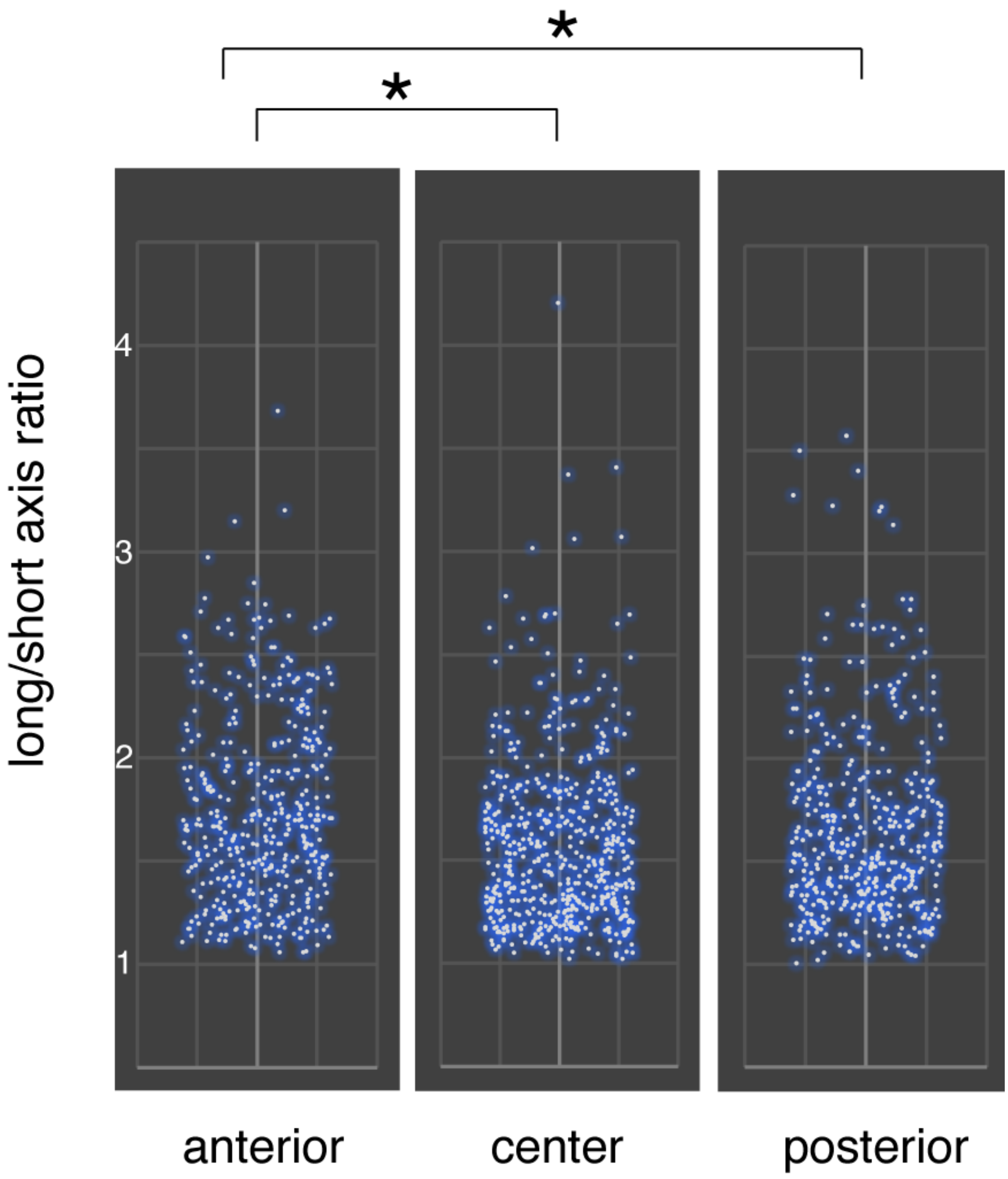


**Extended Data Figure 12. Cell elongation analysis for skate pectoral fins.** The ratio of the long/short axis length of mesenchymal cells in the anterior, center, and posterior pectoral fins was measured and calculated by Fiji (see the Method) and mapped on the graphs. The anterior cells are more oval compared to center and posterior cells. *p* value of t-test between the anterior and center samples is 2.79 x 10^-06 (**) and between the anterior and posterior samples is 0.0195 (*). The numbers of cells investigated in the anterior, center, and posterior fins are 409, 549, and 431, respectively.

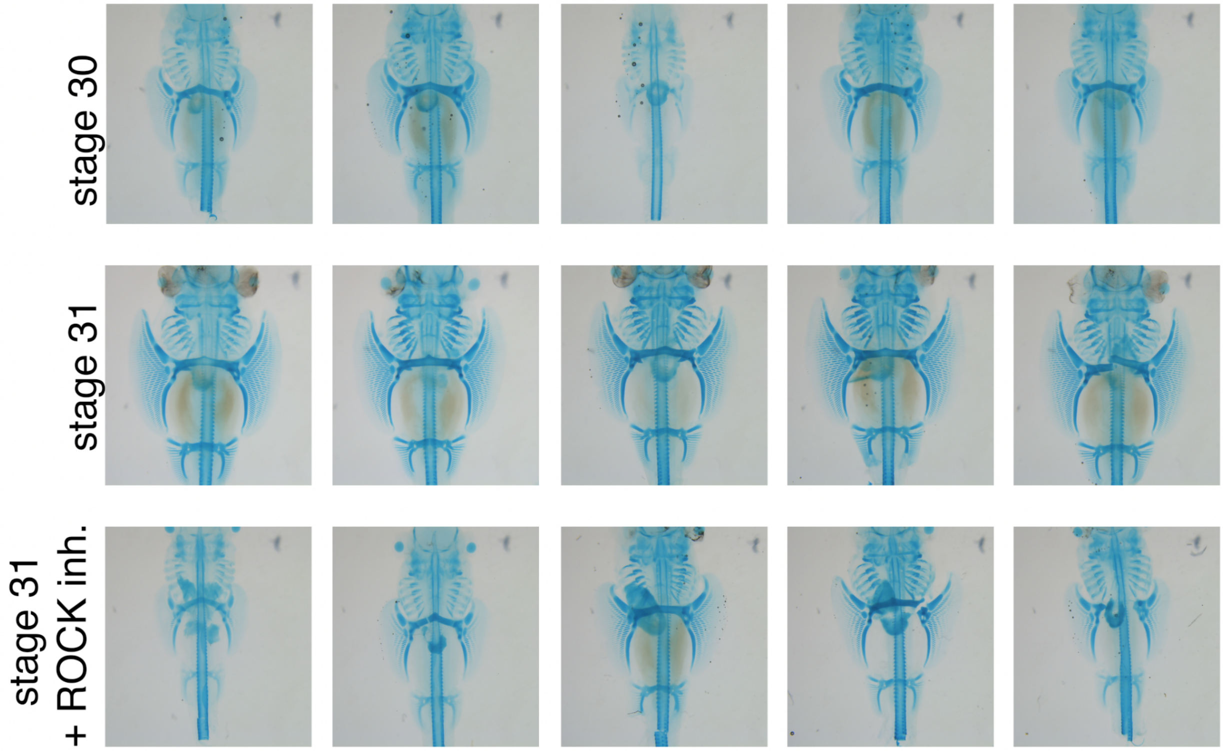


**Extended Data Figure 13. Fin ray development in control and ROCK inhibitor-treated skate embryos.** Cartilages in control (stages 30 and 31) and ROCK inhibitor-treated embryos (stage 31) were examined by Alcian blue staining. Five replicates for each condition are shown. The whole-mount staining showed that anterior fin ray development is affected by ROCK inhibitor-treatment with some variations. The number of fin rays attached to propterygium (pro), mesopterygium (meso), and metapterygium (meta) was counted under a stereomicroscope and statistically analyzed (Fig.4).

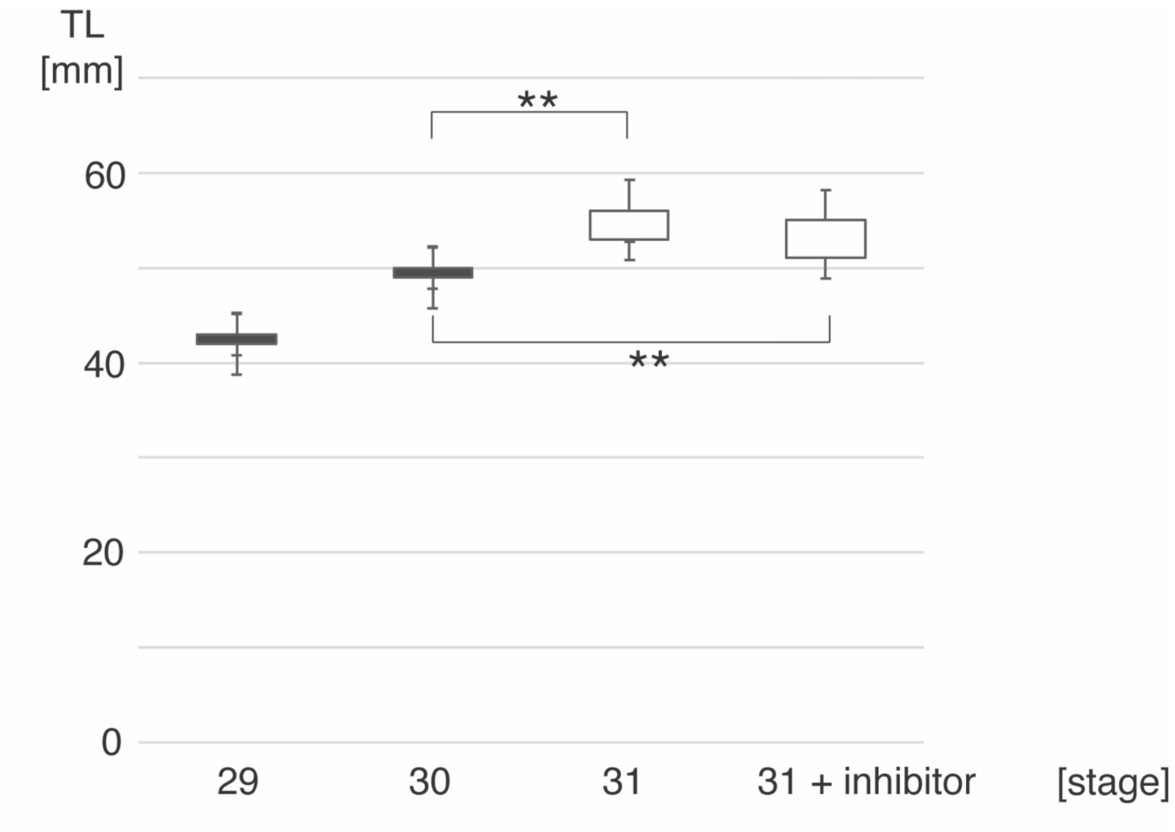


**Extended Data Figure 14. The total body length of control and ROCK-treated skate embryos.** The total body length of control (stages from 29 to 31) and ROCK inhibitor-treated embryos (stage 31). Note that the body length of ROCK inhibitor-treated embryos is longer than stage 30 embryos, indicating that the embryos with the inhibitor normally developed, and the pectoral fin phenotype was not due to the overall defects of body development. Five replicates for each condition were examined.

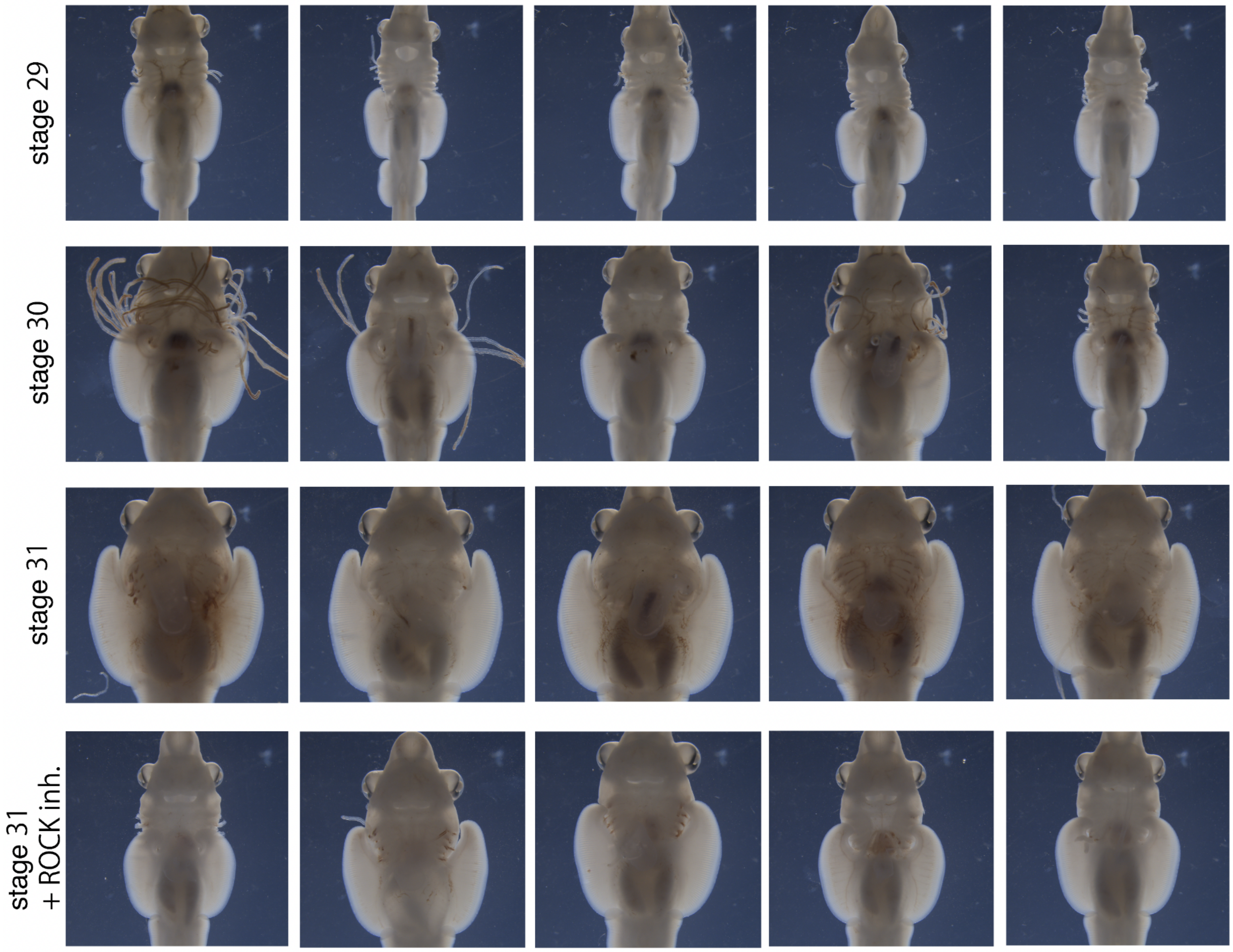


**Extended Data Figure 15. Gross morphology of the pectoral fin in control and ROCK inhibitor-treated embryos.** The pectoral fin develops to the anterior direction from stage 29 to stage 30 and to stage 31. In contrast, the pectoral fin does not grow to the anterior direction in ROCK-inhibitor treated embryos with some phenotypic variations. The external gills were removed for clear visualization of the pectoral fin in some embryos. Five replicates for each stage and condition are shown.

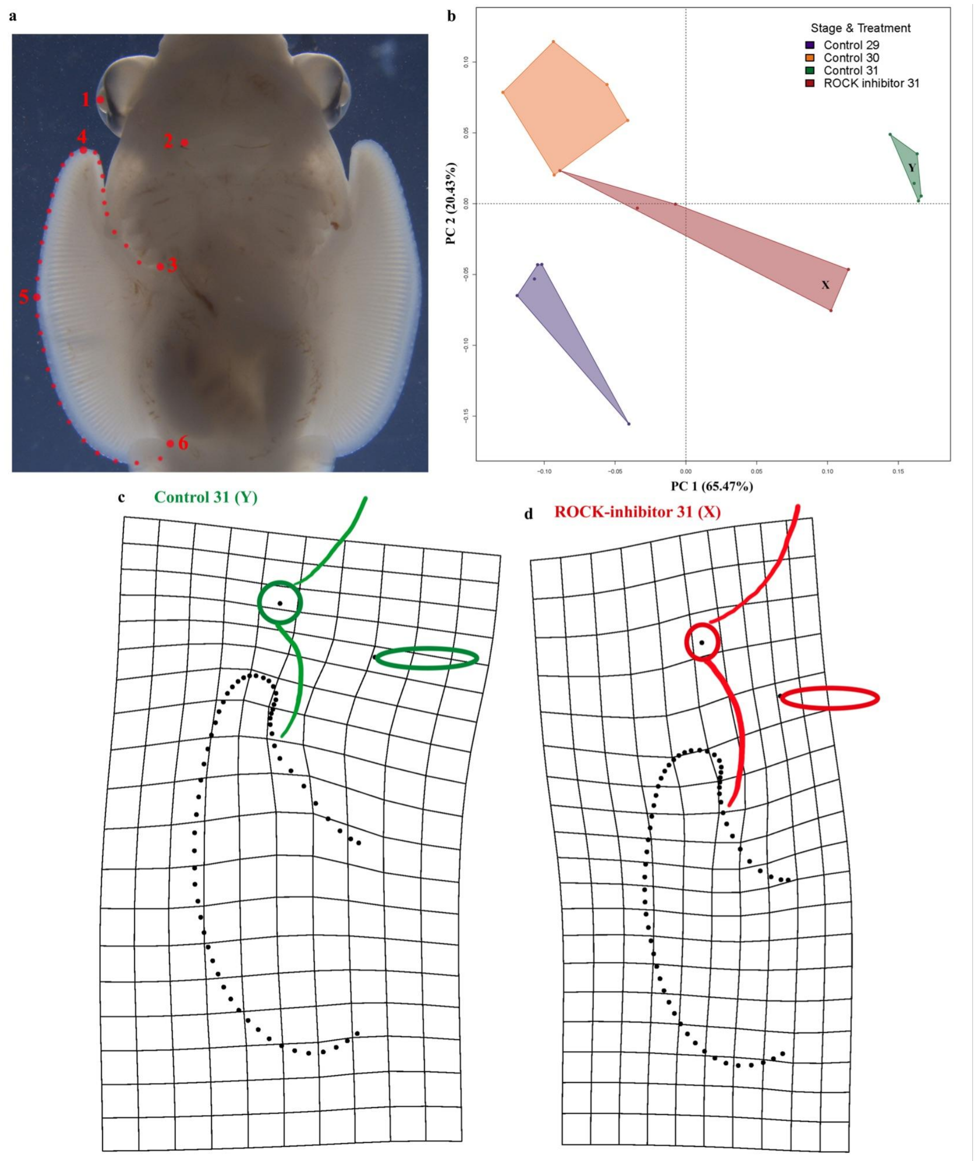


**Extended Data Figure 16.** Geometric morphometric analyses of the inhibition of the PCP pathway using a rho-kinase (ROCK) inhibitor in stage 31 skate embryos. **a**, Schematic of the landmark design used in these analyses, including both landmarks (numbered red points) and semi-landmarks (small red points). **b**, Principal components analysis shows that specimen shapes cluster by treatment and stage. Points X and Y were used to generate the deformation grids showing the shape changes between the area of the PCA plot dominated by control (**c**) and ROCK-inhibited specimens (**d**). Note the inhibition of growth on the anterior region of the pectoral fin in the ROCK-inhibited specimens.

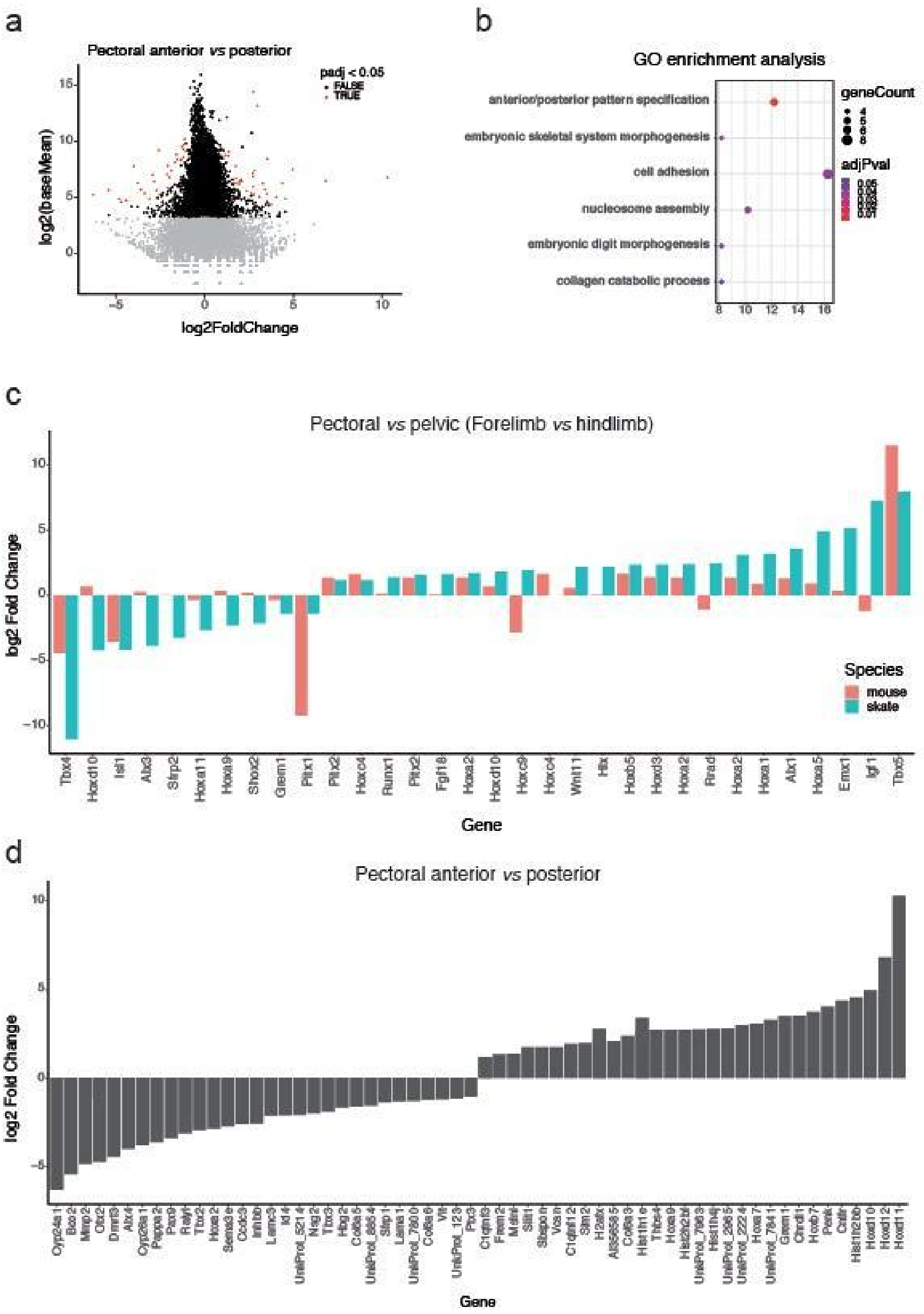


**Extended Data Figure 17. Differential analysis of RNA-seq from anterior and posterior regions of skate pectoral fins. a.** MA plot showing the comparison between anterior and posterior pectoral fins RNA-seq. Statistically significant genes are plotted in red (adjusted p-value < 0.05). **b.** Gene ontology (GO) enrichment analysis of all statistically significant genes. Dot size represents the number of genes in the category, and the color encodes the adjusted p-value. X-axis represents the percentage of genes found from the whole category. **c.** Subset of statistically significant genes when comparing pectoral to. pelvic fins in skate (blue bars), in relation to the same genes when comparing forelimb to hindlimb in mouse (red bars). **d.**

Statistically significant genes from the comparison of anterior *vs.* posterior region of the skate pectoral fin.

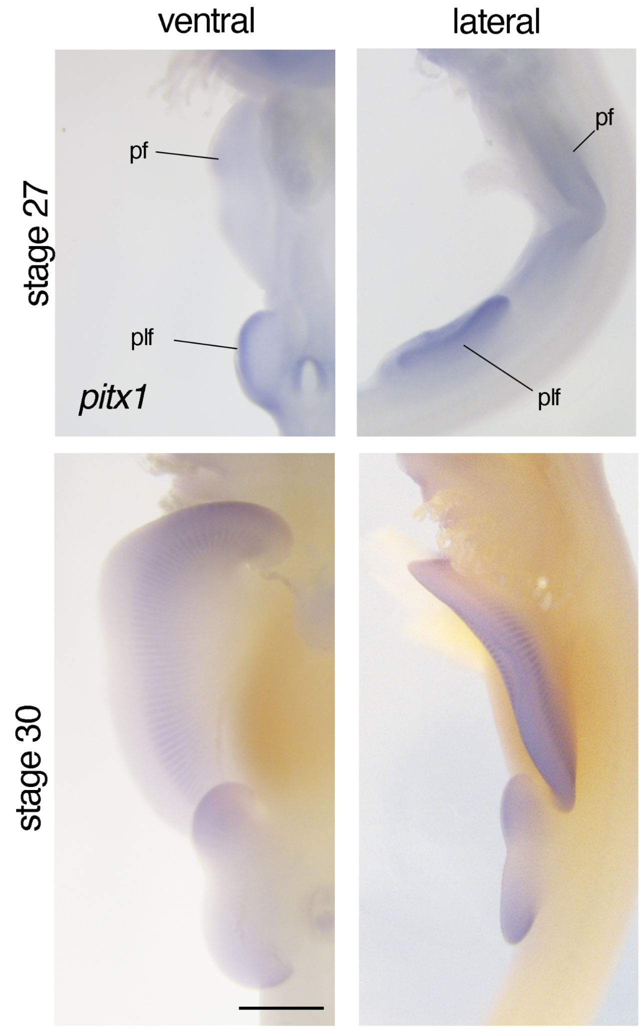


**Extended Data Figure 18. *Pitx1* expression in skate paired fins.**
*Pitx1* is expressed in the pelvic fins (plf) at stage 27. At stage 30, both pectoral (pf) and pelvic fins express *pitx1*. Scale bar= 0.5 mm.

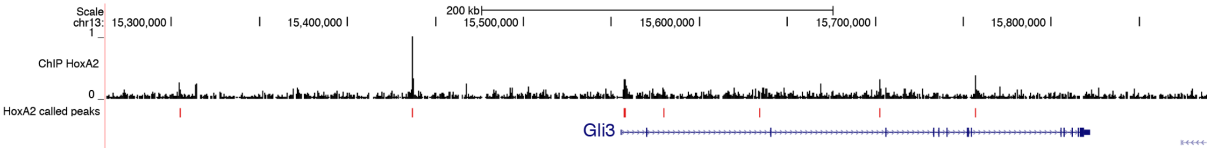


**Extended Data Figure 19. Binding profile of HoxA2 to *Gli3* genomic *locus*.** ChIP-seq experiment in mouse embryonic branchial arches performed in Amin et al. 2015 (PMID: 25640223), which shows the *HoxA2* binding sites around the *Gli3* gene.

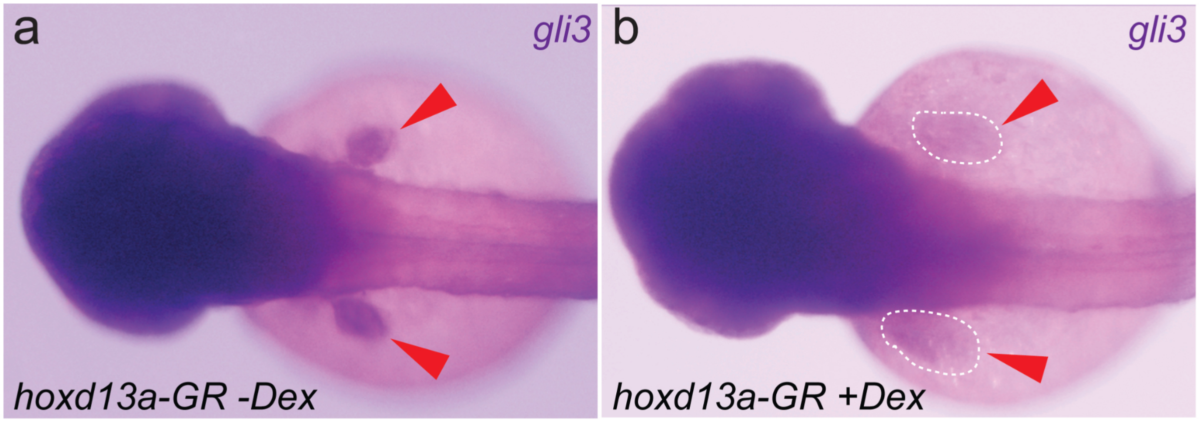


**Extended Data Figure 20. Altered *gli3* expression in *hoxd13a* overexpression mutants.** Whole-mount in situ hybridization of *gli3* in zebrafish embryos carrying a *hoxd13a-GR* overexpression construct. Developing fins are indicated with red arrowheads. **a.** In the absence of dexamethasone, the construct is inactive and the embryos develop normally. **b.** Upon treatment with dexamethasone, *hoxd13a* is activated and causes a reduction of *gli3* expression.

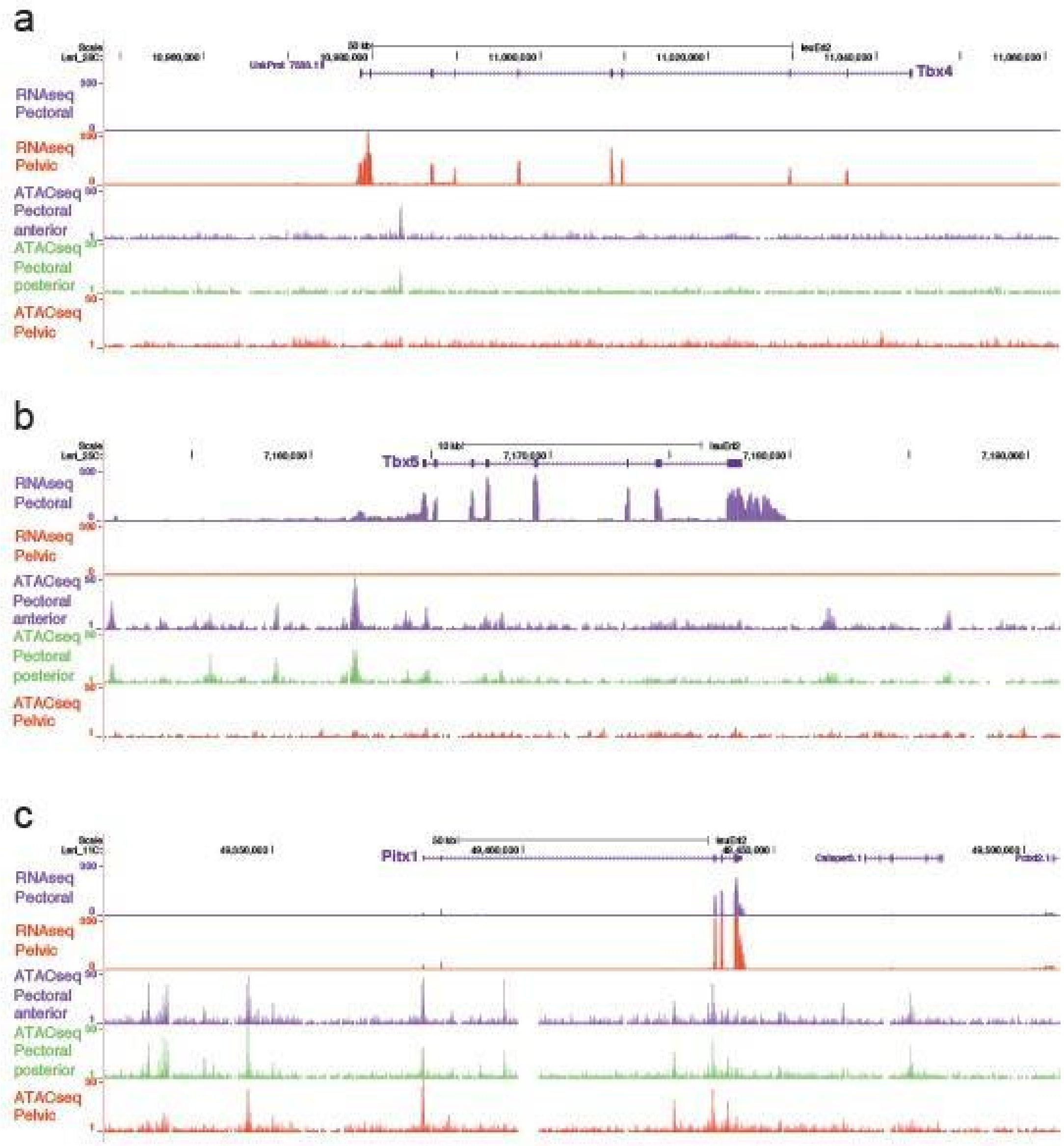


**Extended Data Figure 21. Genomic landscape of *Tbx4*, *Tbx5* and *Pitx1*.** UCSC Browser views surrounding three genes that are key for limb development and anterior-posterior identity. **a, b.** *Tbx4* and *Tbx5* are transcriptionally active in pelvic and pectoral fins, respectively, in a homologous pattern to the expression in mouse, where these genes are active in hindlimb and forelimb, respectively. **c.** Strikingly, *Pitx1* gene is expressed in both pectoral and pelvic fins, while it is only active in hindlimb in other vertebrates.

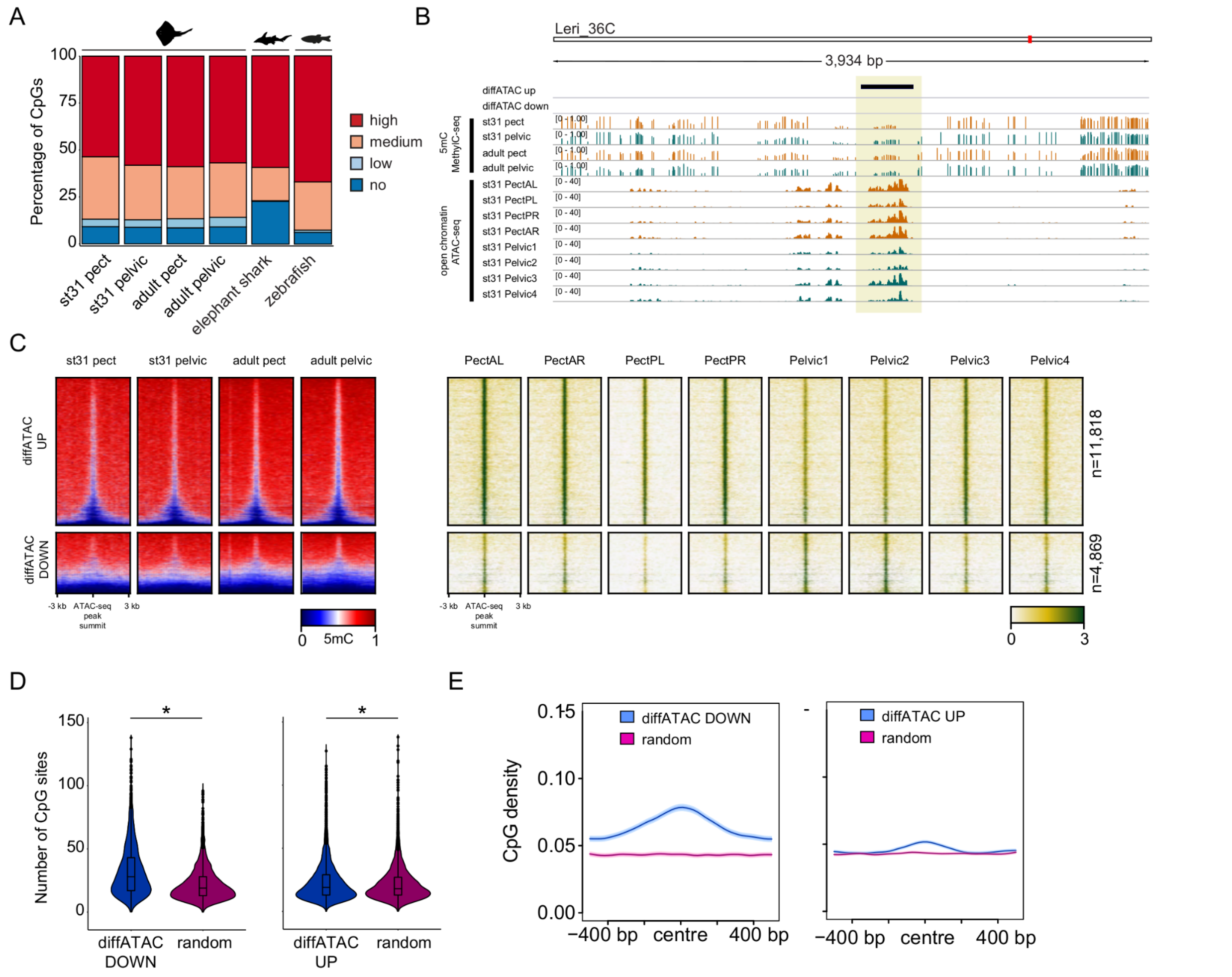


**Extended Data Figure 22. Base resolution DNA methylome profiles of little skate embryonic and adult fins. a.** Stacked bar plots indicating the percentage of methylated CpG sites in little skate (embryonic and adult pectoral and pelvic fins), elephant shark (adult liver) and zebrafish (adult liver) genomes. High, 80-100%; medium, 20-80%; low, >0-20%; no, 0% methylation. **b.** IGV browser tracks depicting DNA methylation (MethylC-seq) profiles at differentially accessible ATAC peaks between pectoral and pelvic fins. **c.** Heatmaps of DNA methylation profiles of differentially accessible ATAC peaks (UP and DOWN) in embryonic and adult pectoral and pelvic fins. Top cluster: ATAC peaks upregulated (diffATAC UP) in pectoral fins. Bottom cluster: ATAC peaks downregulated (diffATAC DOWN) in pectoral fins. **d.** Boxplots depicting the number of CpG sites within differentially accessible ATAC peaks extended +/-500 bp and 1000bp-wide random regions. Wilcoxon rank-sum test was used to compare the number of CpG sites at differentially accessible ATAC regions versus random regions. **e.** Line plots depicting CpG densities of differentially accessible ATAC peaks. The number of CpG sites is calculated in 50bp windows around the ATAC peak centre or around the centre of random regions.

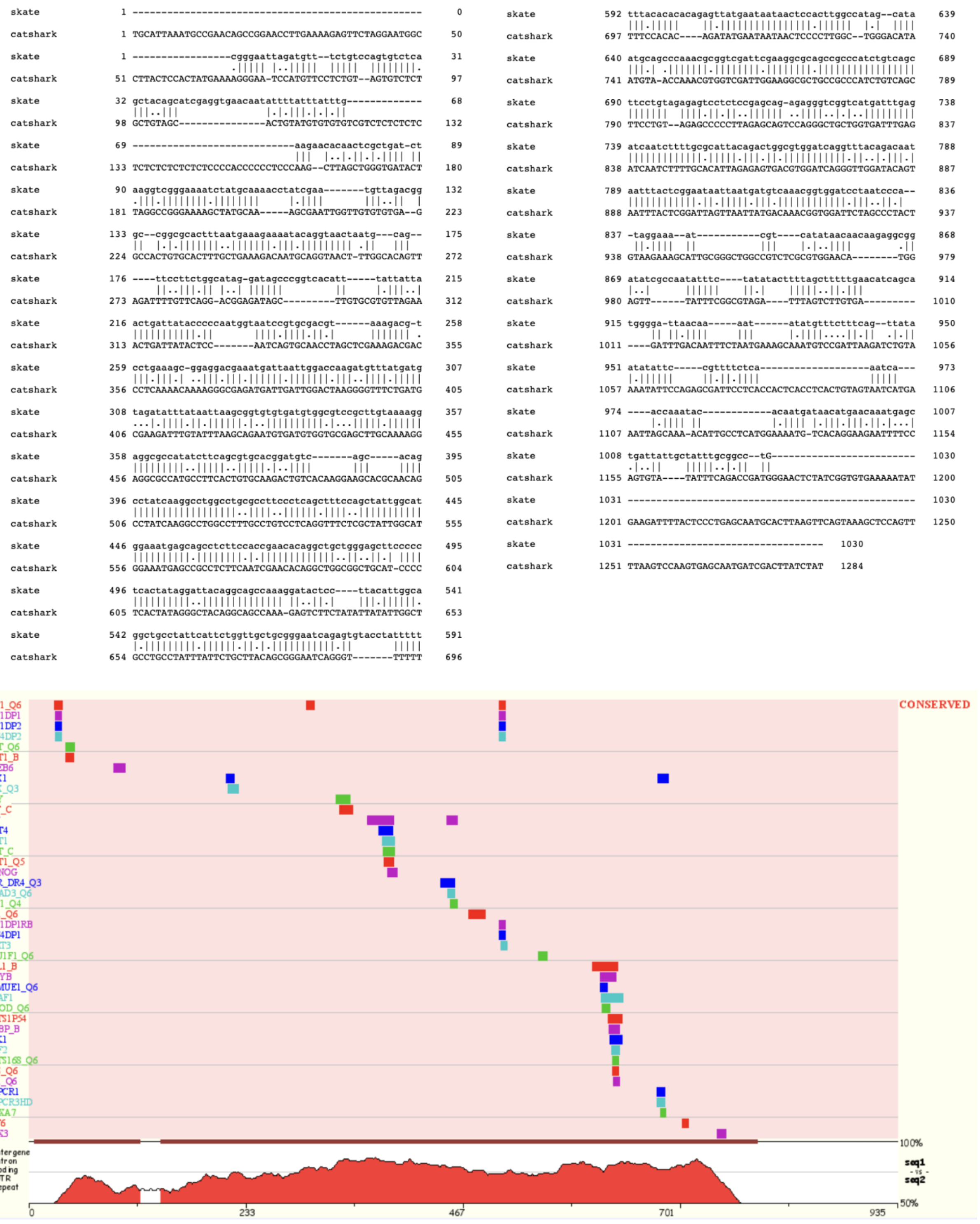


**Extended Data Figure 23. Conservation of sequence and transcription factor binding sites between catshark and skate *HoxA2-1* enhancers.**

Catshark *HoxA2-A1* enhancer (1284 bp) and skate *HoxA2-A1* enhancer (1030 bp) were aligned (top) and conserved transcription factor (TF) binding sites were identified by rVista 2.0 (bottom).

Names and positions of the conserved transcription binding sites are shown by different color codes with sequence conservation peaks (red).
